# A long journey to oyster stem cell markers. *Cg*Sox2 and *Cg*POU2 synergic DNA binding and NODE proteins

**DOI:** 10.1101/2023.09.13.557528

**Authors:** Claude D. Delsert

## Abstract

**Summary statement:** Oyster Sox2 and POU bind in synergy to human Sox-Oct DNA motif, which suggests a role in stem cell maintenance. Sole among the oyster NODE proteins to alter chromatin SalL assembles in large hollow structures that exclude chromatin.

Precursor or stem cells have been identified in mollusk, but determination of differentiation stages is still hampered by lack of markers, notably Oct4, that does not exist among invertebrates. To address this thorny question, oyster Nanog Oct4 DEacetylase proteins were studied, particularly POU proteins, a family that includes vertebrate Oct4.

First, we showed that essential residues of 2 major vertebrate stem cell proteins, Sox2 and Oct4, are conserved through most metazoans which led to experimentally show *in vitro* that recombinant Sox2 protein specifically binds to WT, the mammalian Sox2-Oct4 consensus DNA motif. Furthermore, oyster Sox2 and POU2-3 synergistically bound to WT thus mimicking Sox2 and Oct4 synergy. Altogether, these data suggest that invertebrate POU2 or 3 are Oct4 orthologs. Expression in mammalian cells showed that oyster NODE proteins do not significantly alter chromatin structure except SalL, the homolog of SalL4. Confocal images revealed barrel-shaped hollow structures, orthogonal to cell plane and constituted of aggregates. These bodies were able to fuse or to disassemble/reassemble in fewer but much bigger hollow structures that excluded chromatin.

## Introduction

Bivalve mollusks are among the longest-living animals (Butler et al., 2013) and they are defined as non-senescing species (Finch and Austad, 2001). Indeed, they do not show any age-related decline in physiology or disease resistance. These essential physiological functions depend upon the existence of undifferentiated cells that remained unexplored in mollusks since the precursor work of Cuénot (1891).

More recently, we have shown the existence of undifferentiated cells in oyster such as (i) the adult precursor cells at the origin of hematopoiesis (Jemaà *et al*., 2014) and (ii) the adult primordial germ cells (PGCs) in the gonad very early during reproduction (Cavelier et al., 2017). Nevertheless, the existence of stem cells in mollusk cannot yet be asserted for lack of specific markers and it thus can’t be excluded that precursor cells result from trans-differentiation, a process that, although not preeminent, is known in animals (reviewed in Merrel and Stanger, 2016).

In vertebrates, two transcription factors are essential to stem cells maintenance, Oct4 which expression is restricted to stem cells and Sox2 which in addition is expressed in very early stages of differentiation. The seminal work by Takahashi and Yamanaka (2006) has shown that the ectopic co-expression of Sox2, Oct4, KLF4 and Myc was sufficient to induce mouse cell dedifferentiation into stem-like cells, the induced Pluripotent Stem Cells (iPSC).

Analysis of the genome of an increasing number of invertebrates reveals interesting similarities with vertebrates. Indeed, Önal et al. (2012) showed that the Oct4-activated gene cascade regulating dedifferentiation also exists in planarian despite the fact that the activator, Oct4, does not exist in invertebrates.

In addition, cellular reprogramming of an intestinal epithelial cell into a neuronal cell that naturally occurs in the nematode *Caenorhabditis elegans*, suggests common pathways with the dedifferentiation of iPSC in vertebrates (Kagias et al., 2012). Cellular reprogramming in nematode was shown to depend upon proteins that belong to a Nanog and Oct4-associated deacetylase (NODE)-like complex which enzymatic activity is known to reorganize chromatin through H3 histone deacetylation (Kagias et al., 2012) and methylation (Zuryn et al., 2014). These proteins, namely SEM-4, EGL27, Sox2 and CEH-6, are indeed homologous to the vertebrate SalL4, MTA1, Sox2 and a POU protein, respectively. It is noteworthy that a POU protein is involved in this process, likely replacing Oct4 as POU class V transcription factors do not exist in invertebrates.

In vertebrates, Oct4 and Sox2 play a key role in maintaining the two stem cell-specific potentials, self-renewal and pluripotency. These transcription factors bind two adjacent enhancers, Oct4 and Sox2 DNA-binding elements, to form a complex necessary to regulate the expression of the genes maintaining pluripotency of stem cells, such as Fgf4, Fbxo15, Nanog, Utf1 and Lefty 1, in addition to their own expression (Kuroda et al., 2005).

While Sox2 is expressed in oyster stem or precursor cells (Jemaà et al., 2014; Cavelier et al., 2017), the other members of a putative NODE complex have not been characterized in oyster. The genome of the oyster *Crassostrea gigas* (Zhang et al., 2012; Wang et al., 2019) provided much needed information about putative orthologs of the NODE-like-complex components in oyster.

Because of our long-standing interest in oyster stem/precursor cells, here we report the study of some of the proteins belonging to the NODE complex. While the overall sequence is not well conserved between mammal and invertebrate homologs in general, comparison was performed at the level of functional domains in order to identify the most likely candidates for oyster NODE complex proteins. Functional domains of *Cg*Sox2, *Cg*SalL, *Cg*POUs and *Cg*MTA were examined in search for conservation of key AAs, which are known to be necessary for stem cell maintenance in vertebrates (Tapia et al., 2015). Phylogenetic analysis of these domains revealed that most of the essential AAs in Sox2 and in POU proteins are conserved through metazoans, even in the less evolved animal groups, such as Placozoa, Ctenophora and Porifera. It is noteworthy that Choanozoa, a group of protozoans related to metazoans (King et al., 2008), do not express NODE proteins, suggesting NODE complex is a metazoan innovation.

Purified tagged recombinant oyster POU proteins produced in bacteria were tested *in vitro* through DNA binding assay using an Oct4-Sox2 binding element. Remarkably, *Cg*POU proteins differentially bound to the Oct4-Sox2 DNA element and the DNA binding of *Cg*POU2 or *Cg*POU3 synergistically increased in the presence of *Cg*Sox2.

In addition, inducible stable mammalian cell lines expressing one or two tagged recombinant oyster proteins were selected to study the subcellular localization and potential effects of these proteins on cells, including on chromatin structure. Most proteins were essentially nuclear for *Cg*Sox2, *Cg*POUs and *Cg*SalL as expected for transcription factors. *Cg*SalL formed nuclear bodies that increased in size as the expression time increased. Other proteins, such as *Cg*Lin28, were also present in the cytoplasm in addition to their nuclear location. In particular, *Cg*MTA appeared concentrated in dense foci in the cortical region of the cytoplasm. Altogether, our data show that these oyster proteins behave as their vertebrate homologs and thus have the capability to participate to NODE complexes, hence to participate to oyster stem/precursor cell regulation.

## Results

Homologs of a number of mammalian NODE complex proteins were identified in a number of species belonging to the main metazoan groups (Table S5) using the Conserved Domain Database program (NCBI). Protein domains were examined, in particular for the positional conservation of essential amino acids, as previously determined for Sox2, POUs and SalL4 proteins (Tapia et al., 2015; Pantier et al., 2021).

### Homology of the HMG domain of Sox2

Sox proteins are an animal-specific subclass of the High Mobility Group (HMG) proteins, which is a large family of eukaryotic DNA binding proteins (Stros et al., 2007). Sox2 belongs to the SoxB group of Sox proteins characterized by a well-conserved HMG domain and a little conserved adjacent Soxp domain.

The human and oyster Sox2 HMG differ by only 2 AA substitutions in the 76 AA-long-HMG sequence. Indeed, phylogenic conservation of the HMG sequence through Metazoa is quite remarkable with an overall homology of at least 75% between human and the distant Porifera phylum, which together with the Epitheliozoa constitutes the Metazoa phylum (Simion et al., 2017).

The functionally essential AAs of human Sox2 (K95, R98, M102, R113) (Tapias et al., 2015) are conserved in the human Sox B1 and B2 proteins (Table S1C). More importantly, these AAs are also conserved in the Epitheliozoa phylum that includes most animals at the exception of sponges. Interestingly, these essential AAs are also conserved in certain sponge phyla, including the Homoscleromorpha and some Demospongiae species but they are not present in all Porifera species (Table S1A). It is noteworthy that motifs of the closest HMG protein of *S. rosetta*, a species belonging to Choanozoa, a group of protozoans evolutionary related to metazoans (King et al., 2008), are conserved. But the HMG domain overall homology is weak and it does not contain Sox2 essential AAs (Table S1B).

Soxp is conserved among the jawed vertebrates (Gnathostomata) as illustrated by the homology between human Sox2 and both the bony (*O. latipes*) and cartilaginous (*A. radiata*) fishes. By contrast, Soxp sequence is much less conserved in jawless fish (Lamprey) and its degree of homology decreases as the phylogenetic distance increases among the less evolved organisms (Table S1D).

Conservation of the oyster HMG domain suggests that *Cg*Sox2 might (i) bind to invertebrate DNA sequence related to the vertebrate Oct4-Sox2 DNA elements (Tanimura et al, 2013) (ii) be part of the NODE complex (3i) be involved in cell dedifferentiation (Kagias et al., 2012) or in stem cell maintenance.

### Homology of the functional domains of Oct4

POU transcription factors form a metazoan-specific subclass of the homeo domain proteins, a large family of proteins expressed in all eukaryotes. POUs carry two DNA binding domains, the POU-specific and the homeo domains. POU proteins are divided into 6 subgroups in mammals (POU1-6) while into only 4 or less subgroups in invertebrates. Oct4, a crucial actor of the early development and stem cell maintenance in vertebrates (Leichsenring et al., 2013), belongs to the POU5 subgroup, which remarkably is absent in invertebrates. POU5 likely appeared in an ancestor of Gnathostomata or jawed vertebrates, since the pou5 gene subfamily is missing in the jawless fish (Agnatha) genome (reviewed in Bakhmet and Tomilin, 2022). Indeed, jawless fishes (lamprey, myxine) express only 4 pou genes like invertebrates (Fig. S2B). Based upon their degree of homology with mammals, the oyster POU proteins are categorized as *Cg*POU2, 3, 4 and 6 (Table S2A).

Interestingly, alignment of the POUs domain of human and oyster POUs with human Oct4 shows that 6 of the 7 AAs that are essential for reprogramming iPSCs (Tapia et al., 2015), are conserved only in POU2 and 3. The conserved AAs include I_158_ and D_166_, 2 AAs involved in the formation of the Oct4-Sox2 DNA-binding complex are missing in POU6 (Table S2A, B). Note that K_177_ is very specific of the POU5 subgroup and accordingly, it is conserved only in Gnathostomata (Table S2B). Phylogenetic comparison of POU3, the closest homolog to POU5, shows that the Oct4-essential AAs, except K**_177_**, are conserved throughout the Epitheliozoa (Table S2C).

In Porifera, the situation is not yet clear due to the scarcity of data. Analysis of *O. lobularis* (Homoscleromorpha) genomic sequence provided a partial HMG sequence that contains the Oct4 essential AAs, although it is not known whether this sequence is expressed. Similarly, the essential Oct4 AAs are present in Calcarea and Hexactinellida species. More confusing, in Demospongiae, one species (*O. minuta*) carries the essential AAs while another species (*A. queenslandica*) lacks the Sox2 binding AAs (I_158_, D_166_) (Table S2C).

Alignment of the oyster and human POU homeo domain confirms that *Cg*POU2 and *Cg*POU3 are the closest homologs of human Oct4 while *Cg*POU4 and *Cg*POU6 miss one and three Oct4-essential AAs, respectively (Table S2D).

Conservation of most of the Oct4 essential AAs in the oyster *Cg*POU2 and *Cg*POU3, as is the case for the nematode CEH6 protein (Table S2C, D), suggests that *Cg*POU2 and/or *Cg*POU3 might play the function of Oct4 in the oyster NODE complex.

### Homology of Sal-like and MTA domains

Spalt-like 4 (SalL4), one of the 4 mammalian Sal-Like proteins, maintains vertebrate embryonic stem cell identity and it is required for the development of multiple organs, including limbs. SalL4 is a regulator in the transcriptional network in stem cell maintenance that contains 3 zinc finger regions referred to as ZFC1, 2 and 4. SalL4 mediates its activities through the NODE complex activity (Kagias et al., 2013; Tanimura et al., 2013) but also alternatively, through the direct binding of ZFC4 to AT-rich DNA regions (Watson et al., 2023).

A unique Sal-Like protein exists in mollusk, with 3 conserved zinc finger regions (Table S3) while the rest of the sequence is much less conserved. Interestingly, the two AAs of the ZFC4 domain (T919 and N922), which are essential to SalL4 DNA binding (Watson et al., 2023), are conserved among the bilaterians (Table S3B). Note that SalL proteins are not found in less evolved animals such as the cnidarians, ctenophores, placozoans and sponges for which we aligned unrelated Zinc finger protein sequences that show some degree of homology with SalL4 ZnF4 (Table S3B) but otherwise no homology.

Another component of the NODE complex, the MTA protein, associates with a variety of chromatin remodeling factors which enzymatic activities modulate the plasticity of nucleosomes. The oyster MTA protein contains 5 functional domains (Table S4A), similarly to human MTA1, the largest of three human MTAs: a bromodomain, which recognizes the acetylated lysine on the N-terminal tail of histones, which in turn allows protein-histone association and chromatin remodeling; ELM2 and SANT domains, which interact with DNA binding transcription factors, a GATA binding domain and MTA R1, a MTA1-specific domain (Wang et al., 2008; Kumar and Wang, 2016). Remarkably, the AA sequence of the 5 MTA domains are well conserved between *Hs*MTA1 and *Cg*MTA (Table S4B) while these sequences are not conserved, apart of the SANT domain (Table S4D), in less evolved organisms. Accordingly, while Egl27, the functional orthologue of MTA1 in nematodes (Kagias et al., 2012), does not have a R1 domain and its other domains are only conserved at a lesser degree (Table S4). It is noteworthy that BAH or SANT domains have also been detected in unrelated proteins of Choanozoa although have a lower homology with the animal sequences (Table S4C, D).

### Expression of oyster NODE proteins

To this day, mollusks, in particular marine mollusks, are not easily amenable to genetics or to cell culture. To circumvent this major difficulty hampering protein function studies, we are currently limited to two main possibilities: (i) to show that mollusk proteins can functionally interfere with mammalian proteins *ex vivo* and (ii) to show *in vitro* that mollusk recombinant proteins can interact.

In order to apprehend the regulation of oyster stem or precursor cells, homologs of the afore mentioned proteins, namely *Cg*Sox2, *Cg*MTA, *Cg*SalL and the 4 members of the oyster POU family, were cloned in various eukaryotic and bacterial expression vectors carrying one or more N terminal tags (Materials and Methods).

### *In vitro* DNA binding of the oyster Sox2 and POU proteins

Here, we addressed the intriguing question of which protein(s) if any, can substitute Oct4 in jawless fishes (e.g. lamprey) and in invertebrates, since Oct4 is not encoded in their genome. Interestingly, genetic studies showed that the natural transdifferentiation of an intestinal epithelial cell to a neuron in *C. elegans* is mediated by a deacetylase activity carried out by NODE, a complex that notably contains Sox2, CEH6/POU, SEM4/SalL and Egl27/MTA (Kagias et al., 2012). It is known that mammalian Sox2 and Oct4 proteins synergistically bind to a DNA sequence containing two adjacent short motifs, located in the promoter sequence of genes involved in stem cell maintenance such as Sox2, Oct4, Nanog and a few others (Kuroda et al., 2005; Nakate et al., 2006). The short Sox2-Oct4 binding motifs are slightly variable among these genes and a functional consensus sequence (WT) has been designed (Tanimura et al., 2013).

To test the hypothesis that an oyster POU might bind synergistically with Sox2 to a DNA element *in vitro*, a DNA binding assay was carried out using bacterially expressed *Cg*Sox2 and *Cg*POU recombinant proteins (Materials and Methods). Proteins were purified through glutathione-agarose affinity chromatography, dialyzed and quantified on Coomassie stained SDS-PAGE.

Interestingly, Sox2 binding motifs were identified on the oyster Sox2 gene upstream sequence (data not shown). These motifs were used accompanied by adjacent sequences to design different DNA binding elements, although the adjacent sequences were not close to the Oct4 binding element (data not shown). In addition, the mammalian consensus motif (WT) and its mutated non-functional version (mt) defined by Tanimura et al. (2013) were also tested. Equimolar quantities of recombinant oyster Sox2 or POU proteins were incubated, sole or in combination, with the different biotinylated dsDNA elements and magnetic Streptavidin-agarose beads at 4°C for 1 hr on a rotating wheel. Rinsed beads were boiled in Laemmli buffer and bead supernatant was electrophorezed on 10% SDS-PAGE and transferred to PVDF membrane. Membranes were incubated either with commercial primary anti-Tag antibodies or our in-house immuno-purified anti-*Cg*Sox2 or anti-*Cg*POU2 polyclonal antibodies (Materials and Methods).

Results of an initial experiment revealed using a commercial rabbit anti-GST antibody showed an intense GST-*Cg*Sox2 band when WT DNA was used but only a faint, barely visible band with mt DNA (Fig. 1). Note that GST-*Cg*Sox2 migrated as 55 kDa for a predicted MW of 62 kDa, but Sox2 apparent molecular weight is actually known to be different of its predicted MW. A control using the bacterially expressed C-terminal part of the human p21-Activated Kinase 5, GST-*Hs*PAK5 (Cau *et al*., 2001), showed as expected that it does not bind to WT or to mt DNA or to streptavidin-agarose beads (Fig. 1). GST-*Cg*Sox2 did not bind to mt indicating in addition that it did not artefactually bind to the naïve streptavidin-agarose beads in our experimental conditions.

**Fig. 1.**
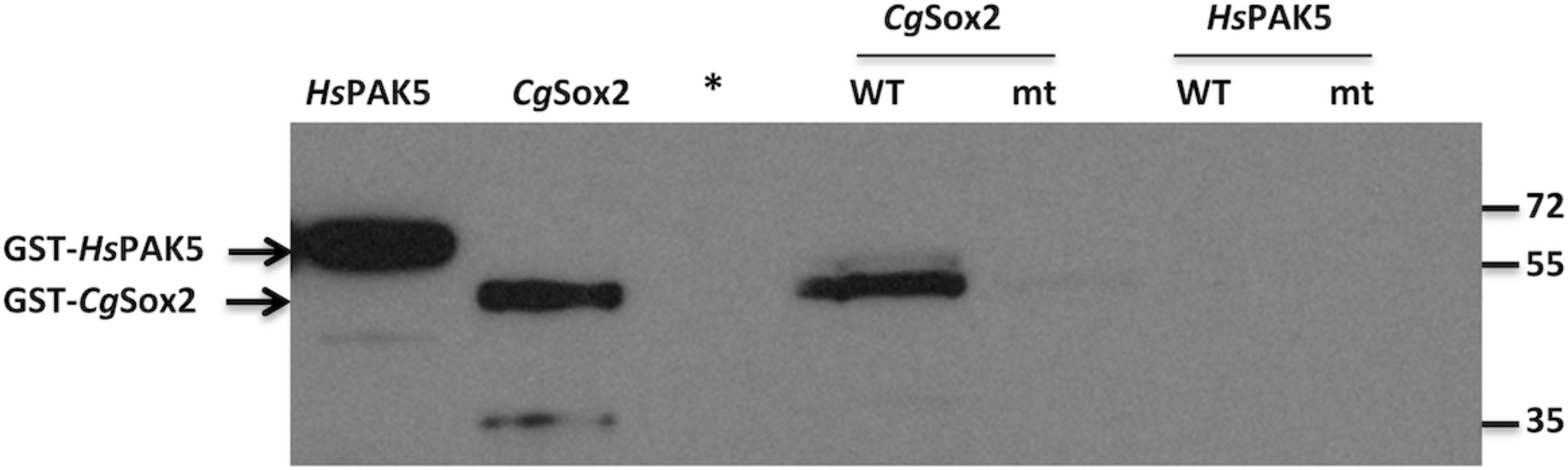
DNA Binding specificity of *Cg*Sox2. Bacterially expressed GST-*Cg*Sox2 and a negative control, GST-Cter*Hs*PAK5, a human signal transduction kinase (Cau et al., 2001) were incubated with WT or mt biotinylated dsDNA before incubation with Streptavidin-agarose beads, rinsed and eluted (Tanimura et al, 2013). Immunoblot of the eluates was revealed using rabbit anti-GST (primary antibody) and anti-rabbit HRP (secondary antibody) and was exposed for autoradiogaphy. Data show that GST-*Cg*Sox2 specifically bound to WT but did not bind to mt DNA while GST-Cter*Hs*PAK5 did not bind to either DNA. * unloaded lane.

It is thus apparent that *Cg*Sox2 strongly and specifically binds to the human Sox2-Oct4 WT DNA element. This unlikely result, considering the phylogenic distance between mollusks and mammals, provided a unique mean to apprehend putative interactions between *Cg*Sox2 and *Cg*POUs.

DNA binding assay was then performed using bacterially expressed His-GST tagged *Cg*POU2, *Cg*POU3, *Cg*POU4 or *Cg*POU6 protein alone or in combination with His-*Cg*Sox2. Immunoblots revealed by a specific anti-*Cg*POU2 antibody showed that *Cg*POU2 DNA binding was specific to WT and that it was strongly increased (>10 fold) in the presence of *Cg*Sox2 (Fig. 2A, upper panel). *Cg*Sox2 binding, revealed by a specific anti-*Cg*Sox2 antibody, showed that Sox2 binding increased in the presence of *Cg*POU2 (Fig. 1A, lower panel). Immunoblots carrying the *Cg*POU3 and *Cg*Sox2 DNA binding eluates were hybridized using a commercial anti-His antibody. Data show that *Cg*POU3 bound WT DNA only in the presence of *Cg*Sox2 while *Cg*Sox2 binding to WT DNA significantly increased in the presence of *Cg*POU3 (Fig. 2B). Similar experiments were performed using *Cg*POU4 and *Cg*Sox2 and revealed using a commercial anti-His (Fig. 2C, upper panel) and our in-house anti-*Cg*Sox2 antibody (Fig. 2C, lower panel) that POU4 bound to WT DNA when Sox2 was present (Fig. 2C). Unlike the above results, *Cg*POU6 bound intensely both the WT and mt DNA (Fig. 2D, left panel), which prompted us to test unrelated DNA motifs, the human SalL4 DNA binding sites 1 and 2. Again *Cg*POU6 bound these two unrelated DNA (Fig. 2D, right panel) and thus no DNA binding specificity could be detected for *Cg*POU6 in our experimental conditions. Thes take home message is that *Cg*POU2 and 3 bind synergistically with *Cg*Sox2 to the Oct4-Sox2 dsDNA motif similarly to human Sox2 and Oct4 (Tanimura et al., 2013). These results should significantly help fishing components of the invertebrate NODE complex and thus should provide stem cell markers to address the thorny question of the existence of stem cells in mollusks.

**Fig. 2.**
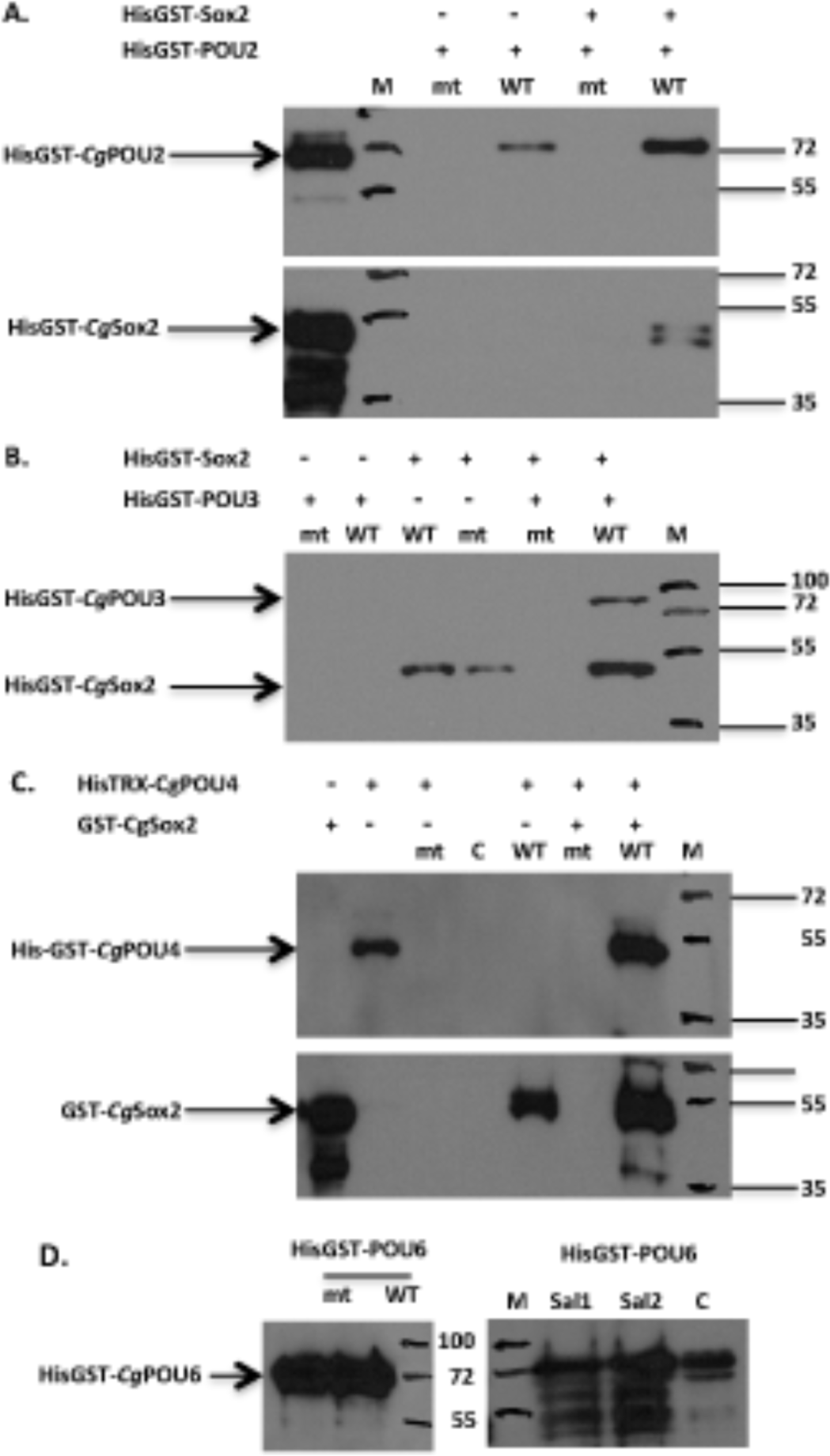
*Cg*Sox2 and *Cg*POU DNA binding. Equal amount of bacterially expressed His and GST tagged proteins were submitted, alone or in combination, to DNA binding using the WT or mt DNA. **A. *Cg*Sox2 and *Cg*POU2 DNA binding.** Eluates of the DNA binding assay were divided in two equal amounts, electrophoresed in parallel on two gels and transferred. First immunoblot was revealed using our in-house rabbit anti-*Cg*POU2 (upper panel). Data show that *Cg*POU2 alone binds to WT but the signal is much more intense (> 10 fold) when *Cg*POU2 is in combination with *Cg*Sox2. Second blot hybridized with in-house rabbit anti-*Cg*Sox2 (lower panel) show a signal (double band due to a protein cleavage) in the presence of WT but not of mt DNA. **B. *Cg*Sox2 and *Cg*POU3 DNA binding.** Immunoblot was hybridized to commercial anti His antibody, which revealed both His-GST-*Cg*POU3 and His-GST-*Cg*Sox2. *Cg*Sox2 bound by itself to WT but much less intensely to mt. *Cg*POU3 did not bind by itself to WT or mt. By contrast, *Cg*POU3 specifically bound to WT in presence of *Cg*Sox2 while *Cg*Sox2 binding was greatly intensified by the presence of *Cg*POU3. **C. *Cg*Sox2 and *Cg*POU4 DNA binding.** Eluates of the DNA binding assay were divided in two, electrophoresed in parallel on two gels and transferred. First immunoblot was revealed using a commercial rabbit anti-His (upper panel). These data show that *Cg*POU4 alone does not bind to WT or to mt. By contrast, the signal is very intense with WT when *Cg*POU4 is in combination with *Cg*Sox2. Second blot was hybridized with in-house rabbit anti-*Cg*Sox2 (lower panel). A signal by the binding of *Cg*POU4 to WT but not to mt. Signal is much stronger when *Cg*POU4 is incubated with *Cg*Sox2. Lane C. incubation of naïve streptavidin-agarose beads with both *Cg*Sox2 and *Cg*POU4. **D. *Cg*POU6 DNA binding.** DNA binding of bacterially expressed His-GST-*Cg*POU6 was revealed by immunoblots using a commercial anti His antibody. An intense signal was produced with WT as well as with mt or with the human SalL4 DNA binding motifs (Sal1 and 2). These results show that *Cg*POU6 did not display binding specificity in our *in vitro* assay. Lane C. Control: native His-GST-*Cg*POU6 protein.

### Oyster NODE proteins in mammalian cell environment

Stable cell lines were established using U2OS cells and doxycycline-inducible pSBTet expression vectors (Kowarz et al., 2015) to express tagged oyster NODE proteins. Cells were treated for immunofluorescence (Materials and Methods) 7 hr after induction. Cells positive for *Cg*Sox2 or *Cg*POU2, 3 or 6 displayed a homogenous fluorescent nuclear signal (Fig. 3) as expected for overexpressed transcription factors. By contrast, MTA although essentially nuclear with a more intense thin peri-nuclear signal, was also present in the cytoplasm (Fig. 4A), in particular as it can be seen at higher magnification, in a few intense foci located in the cortical region (Fig. 4B). Similarly, the signal for the small Lin28 transcription factor was as intense in the cytoplasm as in the nucleus (Fig. 4C).

**Fig. 3.**
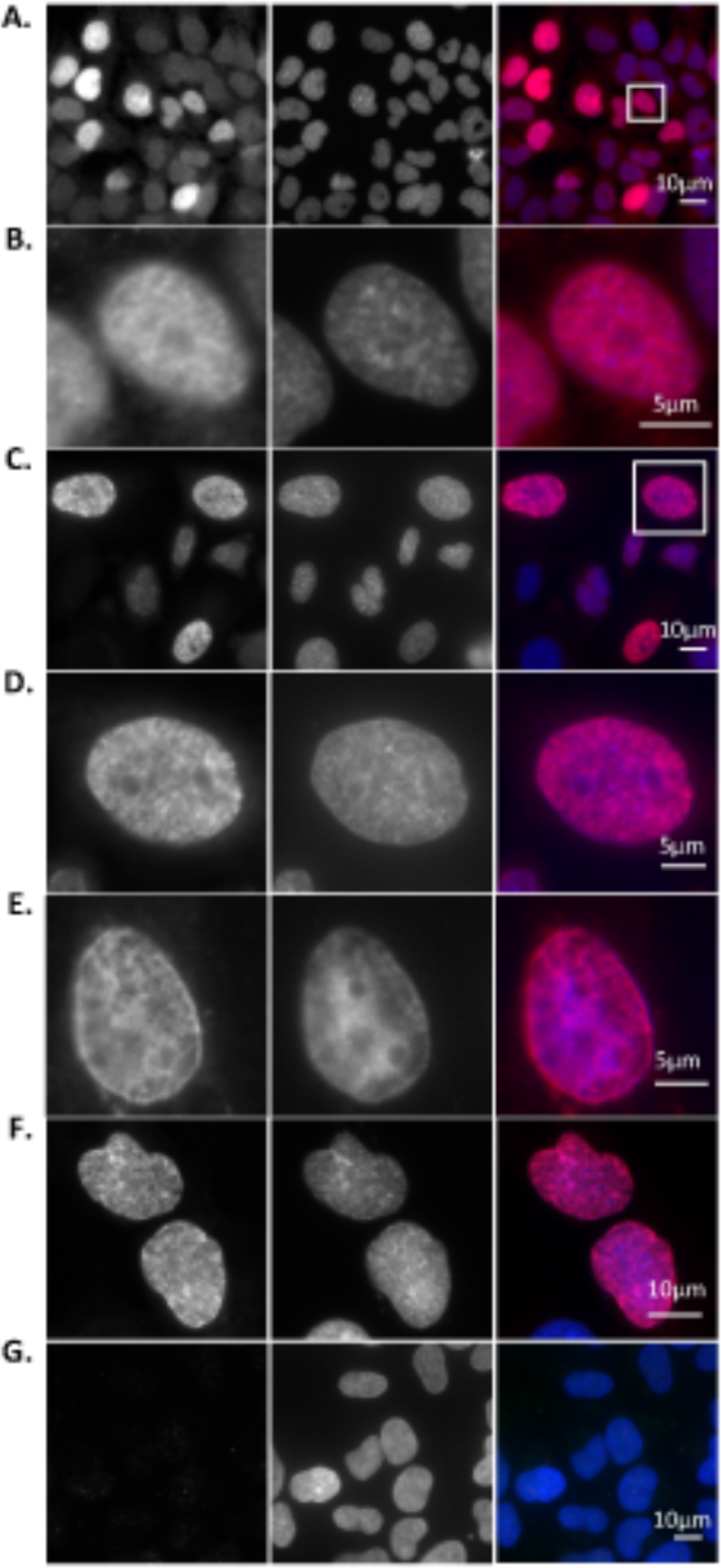
Immunofluorescence of *Cg*Sox2 and *Cg*POUs overexpressed in mammalian cells. U2OS cells were transfected using an inducible expression vector expressing tagged oyster proteins. Cells were induced 16 hrs post-transfection and fixed 16 hrs post-induction. In house specific or commercial anti-tag antibody and fluorescent secondary antibodies were used to reveal protein expression and sub-cellular location. Left panel, oyster protein immunofluorescence; DAPI-stained DNA, middle panel; merged images, right panel. **A. *Cg*Sox2 expression.** In-house specific anti-*Cg*Sox2 antibody revealed the exclusive accumulation of *Cg*Sox2 in the nucleus of transfected cells, indicating that the oyster Sox2 protein is located as its mammalian counterpart. **B. Inset.** Enlargement of the nucleus framed in **A.** shows the diffuse nuclear repartition of *Cg*Sox2 when over-expressed. **C. *Cg*POU2 expression.** In-house specific anti-*Cg*POU2 antibody revealed the exclusive accumulation of *Cg*POU2 in the nucleus of transfected cells, indicating that the oyster POU2 protein is located as its mammalian counterpart. **D. Enlargement of the nucleus framed in C.** IF shows a slightly punctuated signal that suggests local accumulations of *Cg*POU2 in the nucleus. **E. *Cg*POU3 expression.** Commercial anti-HA antibody reveals *Cg*POU3 slightly accumulated in the perinuclear region. Interestingly, *Cg*POU3 also decorates apparent structures of the DAPI-stained chromatin. **F. *Cg*POU6 expression.** IF using the same anti-HA antibody reveals a punctuated fluorescent signal in *Cg*POU6 transfected cells. **G. Negative control**. Fixed U2OS cells were submitted to secondary fluorescent antibody and DAPI staining. In the absence of any primary antibody, IF reveals no signal while DAPI-stained DNA is detected. Scales bars on the lower right side of images.

**Fig. 4.**
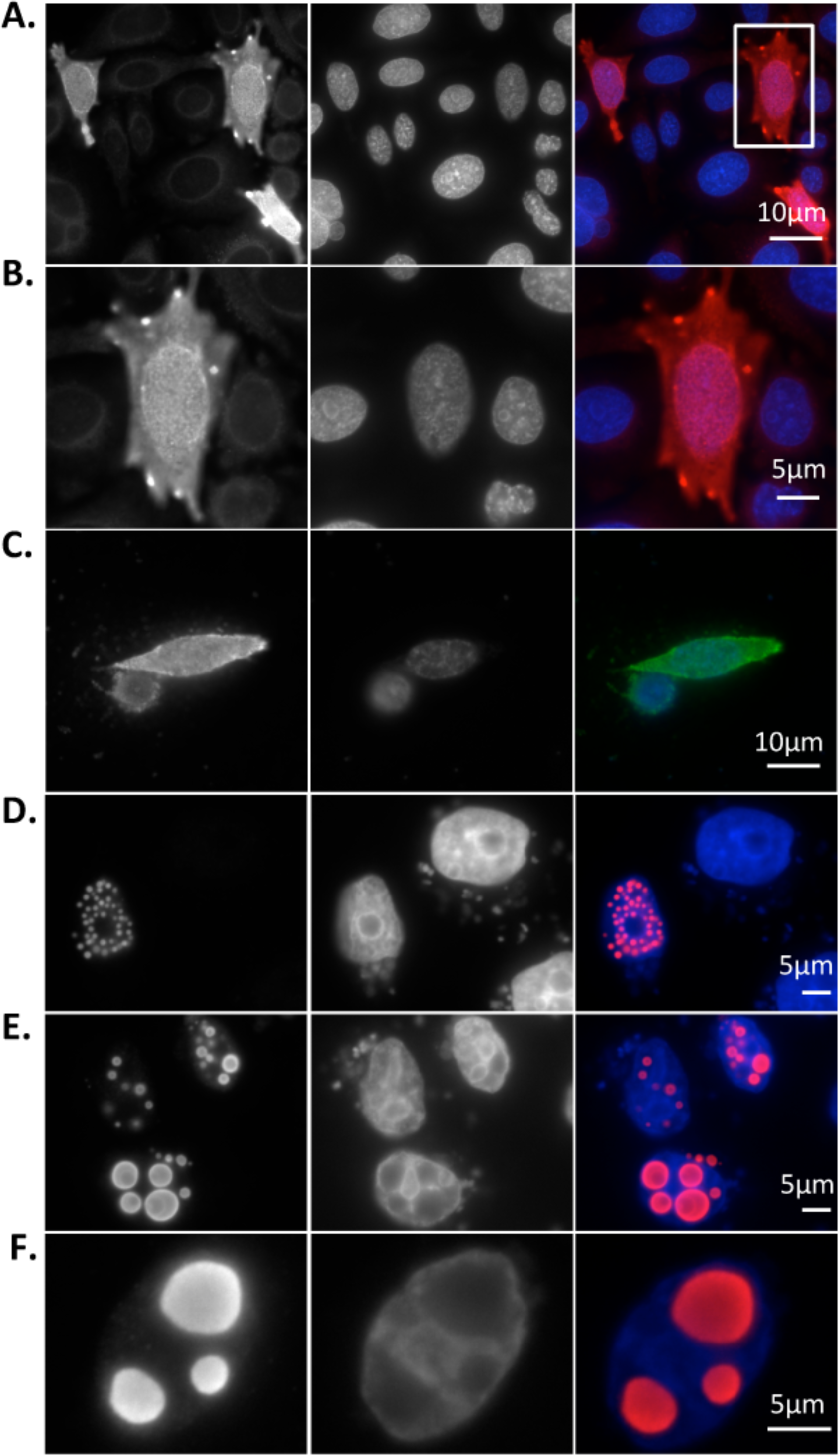
Immunofluorescence of *Cg*SalL, *Cg*MTA and *Cg*Lin28 overexpressed in mammalian cells. U2OS were transfected, Doxycycline induced and fixed for immunostaining as mentioned in Fig. 6 legend. Left panel, oyster protein immunofluorescence; DAPI-stained DNA, middle panel; merged images, right panel. **A. *Cg*SalL IF at 7 hr induction**. IF using commercial anti-Flag antibody revealed small punctuations spread all over the nucleus. **B. *Cg*SalL IF at 16 hr induction**. Less numerous but much larger dots are formed in locations devoid of chromatin. Note that IF signal is more intense in the dots outer edge. **C. Higher magnification of *Cg*Sal at 16 hr induction**. Chromatin exclusion, shown in greater detail, indicates that the oyster SalL protein interacts somehow with the chromatin of mammalian cells. **D. *Cg*MTA IF at 7 hr induction.** *Cg*MTA was detected using a commercial anti-HA antibody in the nucleus but also in the cytoplasm of cells overexpressing this transcription factor. **E. Inset. Enlarged view of *Cg*MTA subcellular location.** Most surprisingly, *Cg*MTA was repeatedly detected as spots in the cytoplasmic cortical area (arrow head). **F. *Cg*Lin28 IF at 7 hr induction.** IF using the anti-HA antibody as above, showed that the overexpressed 28kD transcription factor was not restricted to the nucleus but largely present in the cytoplasm.

By contrast, early expression of *Cg*SalL induced the apparition of dots spread all over the nucleus (Fig. 4D). Dots, rather numerous and small early on (7hrs post-induction)(Fig. 4D), increased in size while diminishing in number in cells with a higher rate of expression (Fig. 4E) or in cells with a longer duration of expression (16 hrs post-induction) (Fig. 4F). It thus appears that dots merge to form quite large bodies as *Cg*SalL protein accumulates. This accumulation led to chromatin exclusion as seen on Fig. 4E and F (DAPI, middle panel).

Confocal microscopy on a stable cell line co-expressing *Cg*Sox2 and *Cg*SalL showed that *Cg*Sox2 was scattered on the whole nucleus while *Cg*SalL formed a limited set of empty and near circular rings corresponding to the dots described above (Fig. 5A). An enlarged view of these rings revealed that they are polygonal rather than circular, with bead-like thickenings (Fig. 5B, left upper panel). Orthogonal projections (FIJI) of a confocal Z stack (Fig. 5B, right upper panel; lower panel) showed that *Cg*SalL rings correspond to sections of barrel-shaped hollow bodies with a length of 1 to 1.1µm and an utmost diameter of 1µm, which is quite reduced at both extremities.

**Fig. 5.**
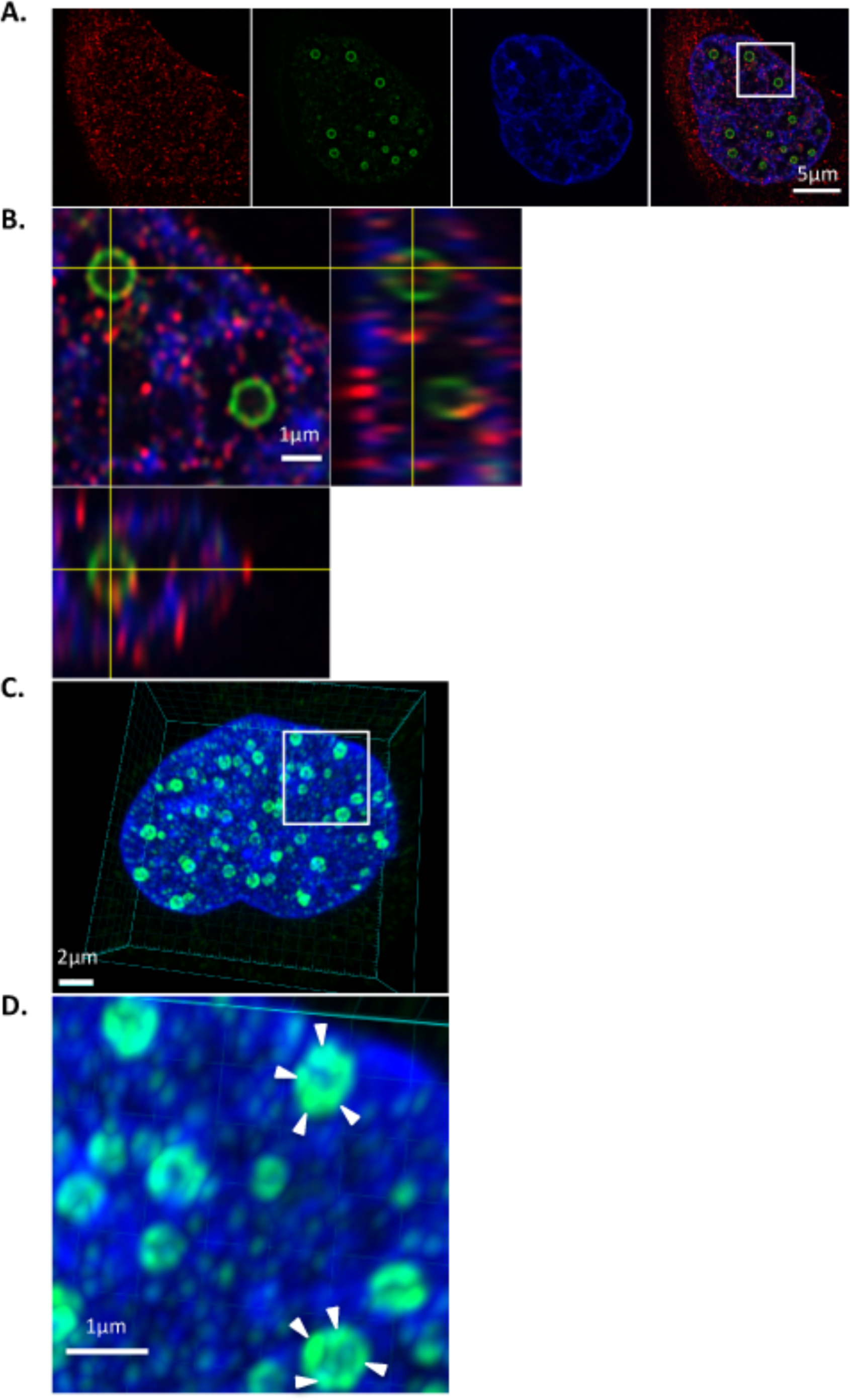
Confocal analysis of *Cg*SalL structures in inducible mammalian cells. Cells stably expressing *Cg*salL and *Cg*Sox2 were induced for 7 hrs using doxycycline, then were fixed and incubated with rabbit anti-*Cg*Sox2 and mouse anti-Flag primary antibodies and stained using anti-rabbit Alexa 555 and anti-mouse Alexa 488 and DAPI. **A. *Cg*SalL nuclear bodies**. Images revealed that *Cg*SalL formed donut-shaped bodies in the nucleus (middle left panel) while *Cg*Sox2 (left panel), red dots evenly spread over the nucleus, did not appear to be associated with these bodies (merge, right panel). **B. 3D analysis of the area framed in A.** Orthogonal projection (FIJI software, NIH, USA) of a middle section of a nuclear stack (Z12 out of 26 sections covering the nuclear depth) (left and lower panels) revealed that *Cg*SalL bodies were actually barrel-shaped hollow volumes of about 0,6 to 1µm of external diameter and 1 to 1.1µm of length. Note that the section appears as an irregular polygon (6 to 8 sides) rather than a circle (upper right panel). Interestingly, these oblong bodies were all perpendicular to the petri dish floor and they were evenly spread in one (sometimes in two) horizontal nuclear plane. **C. More detailed observation of *Cg*SalL nuclear bodies.** Composite 3D image of *Cg*salL nuclear bodies using Imaris (Bitplane) software (Oxford Instruments, UK). **D. Inset.** Higher magnification view suggests that body subunits (arrow heads) contain or are made of *Cg*SalL. *Cg*Sox2 in red (left panel), *Cg*SalL in green (middle left), DAPI stained DNA (middle right), Merge (right panel).

Composite 3D image of *Cg*SalL nuclear bodies using Imaris (Bitplane) software shows that each polygonal ring has a grainy aspect (Fig. 5C). At higher magnification, the grainy rings appear as a string of irregular aggregates of *Cg*SalL proteins (Fig. 5D, arrow heads). Although *Cg*SalL aggregates must be somehow linked to maintain the barrel-shaped structure, they are not fused since space is visible between them. Surprisingly, confocal sections show the same face of these objects, polygonal rings, meaning that these fusiform bodies are all remarkably oriented in the same direction, which is orthogonal to the cell plane.

## Discussion

Preliminary study using the TRAP assay according to Kim et al. (1994), revealed that telomerase activity, a marker for stem/precursor cells, was at its highest in the oyster gonad and gill (unpublished data). This led us to the search for stem or precursor cells in these tissues, a process that led to show their presence and role in hematopoiesis (Jemaà et al., 2014) and gametogenesis (Cavelier et al., 2017) in mollusk. Stem/precursor cells were also observed in other oyster tissues, including mantle, muscle and kidney (unpublished data). Our in-house anti-*Cg*Sox2 antibody was determinant in this characterization although Sox2 is not a marker that allows for the distinction between stem and precursor cells, hence the need for more specific markers. Interestingly, precursor work by Kagias et al. (2012) showed the implication of the NODE complex and Sox2 in promoting cell transdifferentiation in *C. elegans* despite the fact that neither Oct4 nor Nanog exist in invertebrates. Altogether, these evidences led us to search for oyster proteins involved in stem cell maintenance, first through phylogenetic analysis of Sox2 and NODE proteins in metazoans.

Sox2 is a well-conserved protein, notably its HMG box, which in the species we analyzed contains at most only 6 substitutions out of 76 AAs in Epitheliozoa, a group that includes almost all animals and 17 substitutions at most in Porifera (Table S1). Most importantly, Sox2 essential AAs (Tapia et al., 2015) are conserved in all Epitheliozoa and even in some Porifera. The high degree of conservation in the oyster HMG domain suggests that it might bind to vertebrate Sox2 DNA binding motifs. Indeed, DNA motifs homologous to mammalian Sox2 binding sites were found in a 4 kb-sequence encompassing the upstream region and the short coding sequence of the oyster Sox2 gene. Nevertheless, Sox2 motifs were not accompanied by the Oct4 binding motif (unpublished data) as it is the case in mammalian Sox2 and Oct4 promoters.

Kagias et al. (2012) showed that CEH6, the homologue of human POU2, is part of the NODE complex in nematodes, in which it fulfills the function that Oct4 carries out in vertebrates. Search for the essential AAs of the Oct4 POUs domain (Tapia et al., 2012) showed that only POU2 and POU3 retain these AAs at the exception of K177, which is highly specific of the Oct4 protein (Table S2). Remarkably, the POUs domain of POU3 contains the Oct4 essential AAs, again with the exception of K177, in all phylogenetic groups (Table S2C), except in some sponges. Similarly, the POUs and POU_HD_ domains are conserved in *Cg*POU2 and *Cg*POU3, which suggested they might bind to the vertebrate Oct4 binding element.

DNA binding assay was performed essentially as previously described (Tanimura et al., 2013) using His and GST tagged recombinant *Cg*Sox2 or *Cg*POU proteins expressed in bacteria.

Tests using biotinylated double-stranded DNA motifs from the *Cg*Sox2 promoter remained unsuccessful (data not shown), which led us to try the consensus motif for mammalian Sox2 and Oct4 binding (WT) (Tanimura *et al*., 2013). A bit surprisingly, the WT DNA motif proved to bind the recombinant oyster Sox2 while its mutated version (mt) did bind at most only weakly as it does for human Sox2 (Fig. 1).

Moreover, combinations of *Cg*Sox2 and *Cg*POU2 or 3 or 4 induced the synergistic binding of *Cg*Sox2 and *Cg*POU to WT but not to mt (Fig. 2). Concerning POU4, its role in stem cell maintenance is unlikely because its POUs domain contains a 3 AA insert like in all species and it also lacks one of the Oct4 essential AA. On the other hand, *Cg*POU6 lacks the 2 Sox2-binding AAs and we have shown that it strongly and non-specifically binds to unrelated DNAs in our assays (Fig. 2 D).

The DNA protein complex formed by *Cg*Sox2 and *Cg*POU2 or 3 with WT mimics the complex formed by mammalian Sox2 and Oct4 with WT in stem cell extracts (Tanimura et al., 2013). This result is significant as it suggests that *Cg*Sox2 and *Cg*POU2 or 3 could participate to the oyster NODE complex and it follows that it might regulate the maintenance of stem or precursor cells in mollusks. In that regard, it is noteworthy that the first evolutionary occurrence of Oct4 resulted from the duplication of pou2 gene in an ancestor of the cartilaginous fishes (Bakhmet and Tomilin, 2022). Moreover, the POU2 protein from different vertebrates can regulate pluripotency in mammalian stem cells, including in axolotl, in medaka (Tapia et al., 2012) or in chicken (Gold et al., 2014).

To further characterize oyster NODE proteins, the oyster homologues of SEM-4/SalL4, EGL27/MTA1 and Lin28 (Kagias et al., 2012) altogether with *Cg*Sox2 and *Cg*POUs were expressed in mammalian cell cultures through an inducible mammalian vector, as there is no cell culture available for mollusks as for most invertebrates. Immunofluorescence microscopy showed that overexpressed *Cg*Sox2 and *Cg*POU proteins were essentially localized in the nucleus (Fig. 2) as expected for transcription factors. Chromatin of *Cg*POU2 or *Cg*POU6 overexpressing cells appeared slightly punctuated which suggests a certain degree of interaction with mammalian chromatin.

MTA is an important component of the complex since it binds the HDAC1 deacetylase while facilitating contact with the nucleosomes (Millard et al., 2020). *Cg*MTA was localized in the nucleus, notably in the perinuclear region but also in the cytoplasm, in particular in a few spots of the cortical region. These spots appear to be associated with the retraction of the cytoplasmic membrane (Fig. 4B), which suggests the toxicity of the overexpressed MTA. Indeed, longer induction of the expression of *Cg*MTA proved to be lethal (data not shown).

SalL4 is a large protein expressed during development and it is restricted to stem cells in adults (Sweetman and Munsterberg, 2006) where it is bound to both Sox2 and Oct4 proteins (Mallanna et al., 2010; Tanimura et al., 2013). Early during induction (6-7 hrs) *Cg*SalL formed numerous dots in the nucleus while a more intense or longer expression (16 hrs and longer) produced fewer but much bigger nuclear bodies. DAPI-staining showed that formation of these large bodies displaced chromatin (Fig. 4E-F) (see Watson et al., 2023). Sections of confocal image stacks from cells expressing *Cg*Sox2 and *Cg*SalL revealed ring structures that actually corresponded to the dots observed above. Orthogonal projection of a detail of this stack shows that rings are indeed sections of barrel-shaped hollow bodies that are all oriented in the same direction which is orthogonal to the cell culture plane. Analysis of the stacks using the Imaris (Bitplane) software showed at higher magnification that the barrel shaped bodies are made of subunits that form the body walls (Fig. 5D).

The homogeneity of size and structure of *Cg*salL bodies raise a number of questions. *Cg*SalL bodies are made of aggregates organized in homogeneous barrel-shaped hollow structures that would be expected to be stable. Instead, they are apparently labile since they regroup to assemble in less numerous but increasingly larger structures as *Cg*SalL protein concentration increases. This kinetic of disassembly-reassembly to larger bodies is accompanied by chromatin displacement, which is particularly visible when large bodies form (Fig. 4E, F), as shown elsewhere (Watson et al., 2023), must require cargo transport inside the nucleus and thus energy. Regarding the composition of these bodies, it is quite possible that aggregates are made exclusively or mostly of *Cg*SalL since otherwise a tremendously large supply of endogenous protein(s) would be required to match the overexpressed *Cg*SalL protein. Although these homogenous bodies do not represent the functional *Cg*Sal multi-protein complex, their structure can still be of potential interest for understanding their functional structure.

We believe that our results using the biotinylated WT motif pave the way for the characterization of the oyster NODE complex using extracts of fertilized eggs (truly totipotent stem cell) or of larvae at an early stage of development. Identification of the invertebrate Oct4 homologue and other NODE proteins will undoubtedly allow the distinction between precursor and stem cells in invertebrates.

## Materials and Methods

### Cloning and expression vectors

Coding regions were PCR amplified from oyster gill cDNA using specific primers for insertion in Gateway donor vector or primers carrying SfiI sites for directional cloning in inducible pSBTet vectors. Cloning was performed using classical techniques of molecular biology. List of plasmids is summarized in Table S6.

### Recombinant protein and purification

Partial or complete coding sequences were cloned in pGEX or in Gateway expression vectors (Table S6). Accordingly, proteins carried divers tags (GST, His_6_, His_6_-GST, His_6_-TRX) in N terminal position. Bacteria pellets were extracted on ice using sonication in the presence of a mix of protease inhibitors. After centrifugation, clarified protein extract was purified through chromatography on Nickel or on Glutathione Sepharose resins according to manufacturer’s recommendations. Different amounts of purified proteins were electrophorezed on 10% PAGE-SDS and analyzed using Coomassie staining in order to assess both the quality and concentration of each protein batch. Purified proteins were frozen in liquid nitrogen before storage at −70°C.

#### Primary antibodies

In-house rabbit anti-*Cg*Sox2 and anti-*Cg*POU2 sera were prepared as previously described (Jemàa et al., 2014). They were used for immunoblots and immunofluorescence microscopy at dilution of 1/3000 and 1/500, respectively.

### DNA binding assay

The assay was performed essentially as described in Tanimura et al. (2013). Equimolar quantities of recombinant *Cg*Sox2 or *Cg*POU proteins were incubated, sole or in combination, with the WT or mt biotinylated dsDNA elements and magnetic Streptavidin-agarose beads at 4°C for 1 hr on a rotating wheel. Rinsed beads were boiled in Laemmli buffer and supernatant was loaded on 10% SDS-PAGE and transferred to PVDF membrane. Membranes were incubated either with commercial primary anti-Tag antibodies or against in house immuno-purified anti-*Cg*Sox2 or anti-*Cg*POU2 polyclonal antibodies.

Membranes were incubated with Goat anti-rabbit or anti-mouse IgG HRP conjugated (Bio-Rad Corporation, USA) and then with Immobilon Crescendo Western HRP substrate (Millipore, USA) before autoradiography exposition.

#### Stable cell culture selection and immunofluorescence microscopy

Cells were transfected using Doxycycline-inducible pSBTet expression vectors carrying tagged NODE oyster proteins (list of constructs on Table S6) and selected using blasticidine, puromycine or hygromycine. Positive cell cultures were identified through immunoblot analysis and were preserved in liquid nitrogen. Cell cultures at 70 percent confluence were doxycycline-induced for 7 or 16 hrs before fixation using 4% fresh PFA and 0.5% Triton in TBST buffer for 20 mn. Cells were rinsed, incubated using in house-specific anti-*Cg*Sox2 or *Cg*POU2 rabbit polyclonal antibodies or commercial mouse or rabbit anti-tag antibodies and then incubated using Alexa 555- or Alexa 488-labelled fluorescent secondary antibodies to reveal proteins. Coverslips were mounted using Prolong Gold antifade reagent (Life Technologies Corporation, Eugene, USA). Cells were observed using upright **f**luorescence microscopes or a Zeiss LSM880 Airyscan confocal microscope equipped with a ×63 Apo 1.4NA oil objective. Spatial resolution was improved by the use of the airyscan module. Most images are Maximal Intensity Projections (MIP) of 3D Z stacks. 3-dimensional (3D) reconstructed confocal z-stacks were created using Imaris software. Microscopy was performed at the RIO imaging facility in Montpellier (France).

## Acknowledgement

We are indebted to Dr. N. Morin for constant support and to CRBM members for helpful discussions. Special thanks to the Montpellier RIO imaging facility for expert technical assistance (France).

## Data availability

All relevant data can be found within the article and its supplementary information.

## Supplementary information

**Table S1. Phylogenic conservation of the Sox2 HMG and Soxp domains.** Sox genes (SRY-related HMG-box genes) encode a family of transcription factors carrying a conserved HMG box (>50% of homology). **A.** Conservation of Sox HMG box in metazoans. The AAs K95, R98, M102 and R113 (coord. on human Sox2 sequence; highlighted in yellow) of the HMG box are essential to stem cell maintenance (Tapia et al, 2015). They are conserved in Sox-like proteins among all metazoan groups except in certain sponges (Porifera). Note that residue 85 is highly variable. **B.** Conservation in Choanozoa. Alignment of the closest HMG protein of Choanozoa and human Sox2 HMG box indicates a lesser degree of conservation. It notably contains only 1 of the 4 essential AAs of Sox2 HMG box. **C.** Conservation of the HMG domain among the human SoxB proteins. Sox2 essential AAs are conserved in the other human SoxB proteins while Sox2 residue T85 is not (see above). **D.** Phylogenic alignment of Soxp domain. High homology is observed from mammals to fish (both bony and cartilaginous fishes) but not with Agnatha (lamprey) or less evolved organisms except in the Nter sequence adjacent to the HMG box. Coordinates relative to the 1^st^ AA of human HMG; Conserved AAs in red; AAs essential to stem cell maintenance highlighted in yellow.

**Table S2. Phylogenic conservation of POU and homeo domains of POU proteins. A.** Alignment of the POU domain of the 4 oyster POU proteins with human homologs. POU domains of human and oyster POU2 and POU3 are more homologous to Oct4 than POU4 and 6. POU2 and POU3 notably conserve the 2 AAs essential to the binding to Sox2 (red) and the 4 AAs (H154; R157; R186 and K199 in bold letters) necessary to the maintenance of stem cells (Tapia et al, 2012). Note that one lysine (coordinates 177, highlighted in green) is specific of the POUV family, of which Oct4 is member. **B.** Conservation of Oct4 specific AAs in jawless fish. The Oct4 specific lysine (K177, highlighted in green) is found exclusively in Gnathostomes, from human to cartilaginous fish (ray). Indeed, K177 is missing in the early vertebrates (lamprey, Agnathes) while the 6 other essential AAs are conserved. **C.** Phylogenetic conservation of the POU domain in human POU3 homologs. Alignment shows that the essential AAs of the Oct4 POU domain are conserved in POU3, except K177, throughout Epitheliozoa but not in Porifera. Indeed, the essential AAs including the Sox2-POU binding I_21_ and D_26_ (in red) are present in only 3 sponge species belonging to the Homoscleromorpha, Calcarea and Hexactinellida groups. **D.** Conservation of the homeo domain between human and oyster POU proteins. The Oct4 essential AAs are all conserved in POU2 and 3 but not in POU4 and 6 in oyster and in human. Essential AAs highlighted in yellow; Sox2-POU binding AAs (I_158_, D_166_) in red; Oct4-specific lysine (K_177_) highlighted in green.

**Table S3. Phylogenic conservation of Sal-Like Zinc Fingers. A.** Conservation of the human and oyster ZFC1 and ZFC2. Homology is high for ZFC1 with 68% of AA identity (43/63 AAs) (upper alignment) and it is even higher for ZFC2 with 75.8% (47/62) of AA identity (lower alignment). **B**. Phylogenic conservation of ZFC4. This region is well conserved among Protostomia, including Mollusca, with 72% (34/49) of AA identity, in particular with conservation of the 2 AAs essential to stem cells maintenance in position T37 and N41 relative to the first AA of *Hs*ZFC4. Remarkably, a lesser conservation [34% (17/49 AAs) is observed in Plathelminthes and Rotifera. In addition, in Cnidaria and Parazoa this is accompanied by a lack of one or both the T37 or N41 essential AAs. Actually, sequences the closest to ZF4 do not appear to belong to Sal-Like proteins in less evolved animals such as cnidaria, ctenophore, placozoa and Porifera. Identical AAs in red and stem cell essential AAs highlighted in green.

**Table S4. Phylogenic** c**onservation of MTA protein domains. A.** Diagram of the organization of the oyster MTA domains identified using the protein Conserved Domain Database program (NCBI). Note that some domains can be separately found in non-MTA proteins. The oyster MTA domains are similar to human MTA1 while Egl27, the nematode MTA, lacks the R1 region. **B.** Homology between the oyster and human domains. Sequence conservation is markedly high for the BAH, SANT and GATA domains, considering the phylogenetic distance between human and mollusk. **C.** Sequence alignment of metazoan and choanozoan BAH domain. The Nter (upper panel) and Cter (lower panel) sequences of BAH are quite conserved, in particular from mammals to flatworms, while the intervening region separating both domains is quite variable in length and sequence. **C.** SANT domain alignment among metazoans and choanozoans. The SANT domain is a motif of ∼50 amino acids present in proteins involved in chromatin-remodelling and transcription regulation. **D.** Conservation of the oyster and human MTA R1 region. The R1 domain is found in MTA proteins but it is not detected in less evolved organisms, e.g. in *C. elegans*.

**Table S5. Phylogenic position of the organisms cited in this study.** Position of the organisms used in this study provided here for clarity resumes the generally accepted classification. Nevertheless, note that groups we placed at the basis of Epitheliozoa (Ctenophora, Placozoa and Porifera) have currently no consensual taxonomic position. Indeed, any of these 3 metazoan groups can currently be positioned as an out-group of Metazoa depending upon the data analyzed (e.g. see Laumer et al., 2019).

**Table S6. List of the DNA constructs used in this study.**

