## Supplemental Tables for "A long journey to oyster stem cell markers. *Cg*Sox2 and *Cg*POU2 synergic DNA binding and NODE proteins"

### Supplementary Tables

#### Table S1

**A.** (positions 85, 95, 98, 102, 113)

| Sequence | Alignment | Taxon | Species | Accession | Group | Clade |
|---|---|---|---|---|---|---|
| *Hs*Sox2 | RVKRPMNAFMVWSRGQRRKMAQENPKMHNSEISKRLGAEWKLLSETEKRPFIDEAKRLRALHMKEHPDYKYRPRRK | Mammalia | *H. sapiens* | NP_003097.1 | Epitheliozoa | Metazoa |
| *Ol*Sox2 | RVKRPMNAFMVWSRGQRRKMAQENPKMHNSEISKRLGAEWKLLSESEKRPFIDEAKRLRALHMKEHPDYKYRPRRK | Teleostei | *O. latipes* | NP_001265810.1 | Epitheliozoa | Metazoa |
| *Ar*Sox2 | RVKRPMNAFMVWSRGQRRKMAQENPKMHNSEISKRLGAEWKLLSEAEKRPFIDEAKRLRALHMKEHPDYKYRPRRK | Chondrychthian | *A. radiata* | XP_032888050.1 | Epitheliozoa | Metazoa |
| *Bb*Sox2 | RVKRPMNAFMVWSRGQRRKMAQENPKMHNSEISKRLGAEWKLLTEAEKRPFIDEAKRLRALHMKEHPDYKYRPRRK | Cephalochordates | *B. belcheri* | XP_019634859.1 | Epitheliozoa | Metazoa |
| *Aj*Sox2 | RVKRPMNAFMVWSRGQRRKMAQENPKMHNSEISKRLGAEWKLLSESEKRPFIDEAKRLRALHMKEHPDYKYRPRRK | Echinoderm | *A. japonica* | XP_033122142.1 | Epitheliozoa | Metazoa |
| *Cg*Sox2 | RVKRPMNAFMVWSRGQRRKMAQENPKMHNSEISKRLGAEWKLLSEAEKRPFIDEAKRLRAIHMKEHPDYKYRPRRK | Mollusca | *C. gigas* | XP_011455662.1 | Epitheliozoa | Metazoa |
| *Mf*Sox2 | HVKRPMNAFMVWSRGQRRKMAQENPKMHNSEISKILGSEWKKLTEEEKRPFIDEAKRLRALHMKEHPDYKYRPRRK | Annelida | *M. fuliginosus* | QNN94702.1 | Epitheliozoa | Metazoa |
| *Ce*Sox2 | RVKRPMNAFMVWSRGQRKKMALENPKMHNSEISKRLGTEWKMLSEQEKRPFIDEAKRLRAIHMKEHPDYKYRPRRK | Nematoda | *C. elegans* | sp\|Q21305.2 | Epitheliozoa | Metazoa |
| *Of*Sox L | RVKRPMNAFMVWSRERRRRMAQEHPKMHNSEISKRLGAEWKLLSDAEKRPYVDEAKRLRAVHMKDHPDYKYRPRRK | Cnidaria | *O. faveolata* | XP_020630583.1 | Epitheliozoa | Metazoa |
| *Ml*Sox L | RIKRPMNAFMVWSRGQRRKMAQENPKMHNSEISKRLGMDWKMLTEDEKKPFIEEAKRLRALHMKEFPDYKYRPRRK | Ctenophora | *M. leidyi* | AIA23788.1 | Epitheliozoa | Metazoa |
| *Ta*Sox L | RVKRPMNAFMVWSRGQRRKMAQENPKMHNSEISKRLGADWKLLSDADKRPFIDEAKRLRALHMKEHPDYKYRPRRK | Placozoa | *T. adheraens* | RDD46063.1 | Epitheliozoa | Metazoa |
| *Ol*Sox L | RIKRPMNAFMVWSRGQRKKMAHENPKMHNSEISKRLGAAWKLLAETDKRPFMDEAKRLRSVHMKEHPDYKYRPRRK | Homoscleromorpha | *O. lobularis* | OX382163.1 | Porifera | Metazoa |
| *Lc*Sox L | HIKRPMNAFMVWSRDKRKELATQNPKMHNSEISVRLGDEWKSLAEQDKAPFIEEARRLRAQHQADHPDYKYRPRRK | Calcarea | *L. complicata* | AFO66690.1 | Porifera | Metazoa |
| *Sd*Sox L | HIKRPMNAFMVWSKQRRKELAQENPRMHNSELSKRLGSEWKALSDAEKRPYIDEAKKIREQHMIDHPGYRYRPRRK | Demospongiae | *S. domuncula* | CBK62691.1 | Porifera | Metazoa |
| *Ef*Sox L | KVKRPMNAFMVWSRKMRKKIADENPKMHNSEISKRLGAQWKALSDEEKRPYIEEAKRLREAHMKKHPNYKYKPKRK | Demospongiae | *E. fluviatilis* | BAS50201 | Porifera | Metazoa |
| *Aq*Sox L | KVKRPMNAFMVWSRKMRKKIADENPKMHNSEISKRLGTQWKALSEEDKRPFIDEAKRLREAHMKKHPNYKYKPKRK | Demospongiae | *A. queenslandica* | CBK62691.1 | Porifera | Metazoa |
| *Om*Sox L | HIKRPMNAFMVWSKEERKTLSHKNPKMHNSEISKILGTNWKKLSDEDKYPFVEEAKKLRAQHMTDHPGYKYRPKRR | Hexactinellida | *O. minuta* | KAI6651062.1 | Porifera | Metazoa |
| *Om*Sox3 | KVKRPMNAFMVWARDARRKIAQENPKLHNAEISKQLGVSWRELSDEDKSPFRIEAKRIRETHIQKYPQYRYKPRKK | Hexactinellida | *O. minuta* | KAI6654698.1 | Porifera | Metazoa |

**B.**

| Sequence | Alignment | Taxon | Species | Accession | Clade |
|---|---|---|---|---|---|
| *Hs*Sox2 | RVKRPMNAFMVWSRGQRRKMAQENPKMHNSEISKRLGAEWKLLS-ETEKRPFIDEAKRLRALHMKEHPDYKYRPRR | Mammalia | *H. sapiens* | NP_003097.1 | |
| *Sr*HMG | HIKRPMNAFMLWAKDNRAMFIERHPELHNADISRLLGGAWTTRVTDETKRYYAEKAKQVAEEHRRKYPDYKYQPQK | Choanozoa | *S. rosetta* | XP_004996586.1 | Protozoa |

**C.**

| Sequence | Alignment | Accession | Group |
|---|---|---|---|
| *Hs*Sox2 | RVKRPMNAFMVWSRGQRRKMAQENPKMHNSEISKRLGAEWKLLSETEKRPFIDEAKRLRALHMKEHPDYKYRPRRK | NP_003097.1 | SoxB1 |
| *Hs*Sox1 | RVKRPMNAFMVWSRGQRRKMAQENPKMHNSEISKRLGAEWKVMSEAEKRPFIDEAKRLRALHMKEHPDYKYRPRRK | CAA73847.1 | SoxB1 |
| HsSox3 | RVKRPMNAFMVWSRGQRRKMALENPKMHNSEISKRLGADWKLLTDAEKRPFIDEAKRLRAVHMKEYPDYKYRPRRK | NP_005625.2 | SoxB1 |
| *Hs*Sox14 | HIKRPMNAFMVWSRGQRRKMAQENPKMHNSEISKRLGAEWKLLSEAEKRPYIDEAKRLRAQHMKEHPDYKYRPRRK | NP_004180.1 | SoxB2 |
| *Hs*Sox21 | HVKRPMNAFMVWSRAQRRKMAQENPKMHNSEISKRLGAEWKLLTESEKRPFIDEAKRLRAMHMKEHPDYKYRPRRK | NP_009015.1 | SoxB2 |

**D.**

| Sequence | Alignment | Taxon | Species | Accession |
|---|---|---|---|---|
| *Hs*Sox2 | PRRKTKTLMKKDKYTLPGGLLAPGGNSMASGVGVGAGLGAGVNQRMDSYAHMNGWSNGSYSMMQDQLGYPQHPGLNA | Mammalia | *H. sapiens* | NP_003097.1 |
| *Ar*Sox2 | PRRKTKTLMKKDKYTLPGGLLAPGANAMNAGVGVG--VGAAVNQRMDGYAHMNGWTNGGYSMMQEQLGYSQHQGLNA | Chondrychthyan | *A. radiata* | XP_032888050.1 |
| *Hs*Sox2 | PRRKTKTLMKKDKYTLPGGLLAPGGNSMASGVGVGAGLGAGVNQRMDSYAHMNGW-SNGSYSMM--QDQLGYPQHPGLNA | Mammalia | *H. sapiens* | NP_003097.1 |
| *Bb*Sox2 | PRRKTKTLMKKDKYSLPGGLIAANGAPVQRTIATG----------MDQYAHMNGYMSNNAYTHMMHQDQLHAYPHPGISP | Prochordate | *B. belcheri* | XP_019634859.1 |
| *Hs*Sox2 | PRRKTKTLMKKDKYTLPGGLLAPGGNSMASGVGVGAGLGAGVNQRMDSYAHMNGWSNGSYSMMQDQLGYPQHPGLNA | Mammalia | *H. sapiens* | NP_003097.1 |
| *Aj*Sox2 | PRRKTKTLLKKDKYALPG-MLAPGGPGQ-----VQRGVEAMQGYHLNGYAAMNG-VGGYTAMMQEQLHHPYHTQSLS | Echinodermata | *A. Japonica* | \|XP_033122142 |
| *Hs*Sox2 | PRRKTKTLMKKDKYTLPG----GLLAPGGNSMASGVGVGAGLGAGVNQRMDSYAHMNGWSNGSYSMMQDQLGYPQHPGLNA | Mammalia | *H. sapiens* | NP_003097.1 |
| *Cg*Sox2 | PRRKTKTLMKKDKYALPGMPSGAPLQPGREGQMYPMGGYMPNGYPMMHDANAYMNQMGLAGQYGYMPSQQMPGNQMTSGSY | Mollusca | *C. gigas* | XP_011455662.1 |
| *Hs*Sox2 | PRRKTKTLMKKD-KYTLPGGLLAPGGNSMASGVGVGAGLGAGVNQRMDSYAHMNGWSNGSYSMMQDQL--GYPQHPGLNA | Mammalia | *H. sapiens* | NP_003097.1 |
| *Of*Sox2 | PRRKSKTLLKKDNKYTLSMLG-AQGGPPVQRSVAQNPAEHFAQMNGFSAYGPITAYSQMNVNDPYSTMYAGHPLSPHTPT | Cnidaria | *O. faveolata* | XP_020630583.1 |

#### Table S2

**A.** (positions 138, 154, 157, 158, 166, 177, 186, 199, 212)

| Sequence | Alignment | Organism | Species | Accession |
|---|---|---|---|---|
| *Hs*Oct4 | DIKALQKELEQFAKLLKQKRITLGYTQADVGLTLGVLF---GKVFSQTTICRFEALQLSFKNMCKLRPLLQKWVEEAD | human | *H. sapiens* | NP_001272916.1 |
| *Cg*POU3 | DDAPSSDDLEAFAKQFKQRRIKLGFTQADVGLALGTLY---GNVFSQTTICRFEALQLSFKNMCKLKPLLQKWLEEAD | oyster | *C. gigas* | EKC31695.1 |
| *Hs*POU3 | EDTPTSDDLEQFAKQFKQRRIKLGFTQADVGLALGTLY---GNVFSQTTICRFEALQLSFKNMCKLKPLLNKWLEEAD | human | *H. sapiens* | XP_034335340.1 |
| *Cg*POU2 | ENIDLEELLEQFAKTFKRRRIELGFTQGDVGLAMGKLY---GNDFSQTTISRFEALNLSFKNMCKLKPLLQKWLKDAD | oyster | *C. gigas* | EKC20170.1 |
| *Hs*POU2 | EEPSDLEELEQFARTFKQRRIKLGFTQGDVGLAMGKLY---GNDFSQTTISRFEALNLSFKNMCKLKPLLEKWLNDAE | human | *H. sapiens* | XP_011541043.1 |
| *Cg*POU4 | TECDPRELLEAFAERFKQRRIKLGVTQADVGAALANLKLPGVGSLSQSTICRFESLTLSHNNMIALKPILQAWLEEAE | oyster | *C. gigas* | XP_011456596.1 |
| *Hs*POU4 | DVESDPRELEAFAERFKQRRIKLGVTQADVGAALANLKIPGVGSLSQSTICRFESLTLSHNNMIALKPVLQAWLEEAE | human | *H. sapiens* | NP_004566.2 |
| *Cg*POU6 | VDGINLDEIKEFAKQFKIRRLSLGLTQTQVGQALSATE---GPAYSQSAICRFEKLDITPKSAQKIKPVLERWMQEAE | oyster | *C. gigas* | EKC42925.1 |
| *Hs*POU6 | EDGINLEEIREFAKNFKIRRLSLGLTQTQVGQALTATE---GPAYSQSAICRFEKLDITPKSAQKLKPVLEKWLNEAE | human | *H. sapiens* | NP_001357888.1 |

**B.**

| Sequence | Alignment | Organism | Species | Accession | Group |
|---|---|---|---|---|---|
| *Mm*Oct4 | DMKALQKELEQFAKLLKQKRITLGYTQADVGLTLGVLF---GKVFSQTTICRFEALQLSLKNMCKLRPLLEKWVEEAD | mouse | *M. musculus* | NP038661.2 | Gnathostoma |
| *Hs*Oct4 | DIKALQKELEQFAKLLKQKRITLGYTQADVGLTLGVLF---GKVFSQTTICRFEALQLSFKNMCKLRPLLQKWVEEAD | human | *H. sapiens* | NP_001272916.1 | Gnathostoma |
| *Ol*POU5 | EENLSTEELEQFAKELKHKRITLGFTQADVGLALGNLY---GKMFSQTTICRFEALQLSFKNMCKLKPLLQRWLDEAE | bony fish | *O. latipes* | NP_001098339 | Gnathostoma |
| *Ar*POU5 | EEEYPTK-MEQFAKELKHKRITLGFTQADVGLALGNLY---GKMFSQTTICRFEALQLSFKNMCKLKPILQRWLNDAE | ray | *A. radiata* | XP_032886709.1 | Gnathostoma |
| *Pm*POU3 | PAPPTSDDLEQFAKEFKQRRIKLGFTQADVGLALGTLY---GNVFSQTTICRFEALQLSFKNMCKLKPLLSKWLEEAD | lamprey | *P. marinus* | XP_032828582.1 | Agnatha |
| *Pm*POU2 | EDPRDLEELEVFAKMFKQRRIKLGFTQGDVGLAMGKLY---GNDFSQTTISRFEALNLSFKNMCKLKPLLEKWLNDTE | lamprey | *P. marinus* | XP_032821336.1 | Agnatha |
| *Pm*POU4 | DTEADPRELEAFAERFKQRRIKLGVTQADVGSALGNLKIPGVGSLSQSTICRFESLTLSHNNMIALKPILQAWLEEAE | lamprey | *P. marinus* | XP_032811279.1 | Agnatha |
| *Pm*POU6 | VDGVNLEEMREFARAFKIRRLSLGLTQTQVGQALSAAE---GPAYSQSAICRFEKLDITPKSAQKIKPVLERWMEEAE | lamprey | *P. marinus* | XP_032807447.1 | Agnatha |

**C.**

| Sequence | Alignment | Organism | Species | Accession | Taxon | Group | Clade |
|---|---|---|---|---|---|---|---|
| *Hs*POU3 | EDAPSSDDLEQFAKQFKQRRIKLGFTQADVGLALGTLY---GNVFSQTTICRFEALQLSFKNMCKLKPLLNKWLEETD | human | *H. sapiens* | XP_034335340.1 | Mammalia | Epitheliozoa | Metazoa |
| *Ar*POU3 | EETPTSDELEQFAKLFKQRRIKLGFTQADVGLALGTLY---GNVFSQTTICRFEALQLSFKNMCKLKPLLNKWLEEAD | sting ray | *A. radiata* | XP_032886709.1 | Chondrychthyan | Epitheliozoa | Metazoa |
| *Pm*POU3 | PAPPTSDDLEQFAKEFKQRRIKLGFTQADVGLALGTLY---GNVFSQTTICRFEALQLSFKNMCKLKPLLSKWLEEAD | lamprey | *P. marinus* | XP_032828582.1 | Agnatha | Epitheliozoa | Metazoa |
| *Bb*POU3 | DDTPTSDDLEQFAKQFKQRRIKLGFTQADVGLALGTLY---GNVFSQTTICRFEALQLSFKNMCKLKPLLQKWLEEAD | lancelet | *B. belcheri* | XP_019614359.1 | Cephalochordata | Epitheliozoa | Metazoa |
| *Aj*POU3 | EEAPTSDDLEQFAKMFKQRRIKLGFTQADVGLALGTLY---GNVFSQTTICRFEALQLSFKNMCKLKPLLAKWLEEAD | Annessia | *A. japonica* | XP_033095503.1 | Echinodermata | Epitheliozoa | Metazoa |
| *Cg*POU3 | DDAPSSDDLEAFAKQFKQRRIKLGFTQADVGLALGTLY---GNVFSQTTICRFEALQLSFKNMCKLKPLLQKWLEEAD | oyster | *C. gigas* | EKC31695.1 | Mollusca | Epitheliozoa | Metazoa |
| *Sm*POU3 | NQKISTDELEQFAKIFKQRRIKLGFTQADVGLALGNLY---GSVFSQTTICRFEALQLSFKNMCKLRPLLGKWLQEAD | Planaria | *S. mediterranea* | JN837708.1 | Plathelminthes | Epitheliozoa | Metazoa |
| Ceh18 | D-RIDMNELEAFAQTFKKQRIKFGFTQGDVGVALGKRY---GTDFSQTTISRFEALNLSFKNMCKLRPLLKEWLADVE | nematode | *C. elegans* | NP_00139107 | Nemathelminthes | Epitheliozoa | Metazoa |
| *Nv*POU | DDTPTSDDLEQFAKQFKQRRIKLGFTQADVGLALGTLY---GNVFSQTTICRFEALQLSFKNMCKLKPLLQKWLEEAD | hexacorallia | *N. vectensis* | AOP31985.1 | Cnidaria | Epitheliozoa | Metazoa |
| *He*POU3 | SAAITTDDLEAFAKEFKRKRIKLGFTQSDVGLGLGSLY---GNIFSQTTICRFEALQLSFKNMCKLQPLLVKWLDEMD | Hydrozoa | *H. echinata* | AEG66930.1 | Cnidaria | Epitheliozoa | Metazoa |
| *Ar*POU | DTSPSSDDLEQFAKEFKQRRIKLGFTQADVGLALGTLY---GNVFSQTTICRFEALQLSFKNMCKLKPLLAKWLEEAD | Scyphozoa | *A. aurita* | AIC75315.1 | Cnidaria | Epitheliozoa | Metazoa |
| *Am*POU | DDTPTSDDLEQFAKEFKQRRIKLGFTQADVGLALGTLY---GNVFSQTTICRFEALQLSFKNMCKLKPLLAKWLEEAD | Anthozoa | *Acropora* | ABD97868.1 | Cnidaria | Epitheliozoa | Metazoa |
| TriPOU | EDTPTSDDLEQFAKQFKQRRIKLGYTQADVGLALGSLY---GNVFSQTTICRFEALQLSFKNMCKLKPLLNKWLEQAD | Placozoa | *T. adheraens* | RDD45971.1 | Placozoa | Epitheliozoa | Metazoa |
| *Olo*POU | -------ELEQFAKMFKQKRIKLGFTQADVGQAL-TDY---GNPFSQTTICRFEALQLSFKNMVKLRPLLHRWLEDAE | Homosclero. | *O. lobularis* | OX382160.1 | Porifera |  | Metazoa |
| *Sc*POU | KPTPSADHLRAFAKQLKLKRIRKGFTQHEVGRDLGLMY---HQNFSQTTICRFEAQQLSHKNMCKLYAILKRWSEDVD | Calcarea | *S. ciliatum* | CAH0790952 | Porifera | | Metazoa |
| *Om*POU | LKEPTQPEMEGFAKEFKQKRISFGFTQADVGNELGKLY---GNAFSQTTICRFEAMQLSFRNMCKLRPILLHWLNEVD | Hexactinellida | *O. minuta* | KAI6649082.1 | Porifera | | Metazoa |
| *Aq*POU | E-SPEVKNLEAFAVSFKSRRVKLGYTQTNVGQALASVH---GTNFSQTTICRFENLQLSYKNAQKLKPILEKWLEEAE | Demosponge | *A. queensland* | XP01141053 | Porifera | | Metazoa |
| *Hd*POU | PESPEVKDLESFATSFKTKRLKLGFTQTQVGQALAEIN---GTDFSQTTICRFENLQLSFKNAQKLKPVLEKWLEQAE | Demosponge | *H. dujardinii* | MT272763 | Porifera | | Metazoa |

**D.**

| Sequence | Alignment | Organism | Species | Accession |
|---|---|---|---|---|
| *Ol*POUL | RRTTISIPGKEALEQHFLRQQKPSPKEITKIANYLQLEKEVVRVWFCNRRQREKR | Sponge | *O. lobularis* | OX382160.1 |
| *Hs*Oct4 | KRTSIENRVRGNLENLFLQCPKPTLQQISHIAQQLGLEKDVVRVWFCNRRQKGKR | human | *H. sapiens* | NP_001272916.1 |
| *Cg*POU3 | KRTSIEVTVKGALENHFMKQPKPAAQEISQLAEQLQLEKEVVRVWFCNRRQKEKR | oyster | *C. gigas* | EKC31695.1 |
| *Hs*POU3 | KRTSIEVSVKGALESHFLKCPKPSAQEITSLADSLQLEKEVVRVWFCNRRQKEKR | human | *H. sapiens* | XP_034335340.1 |
| *Cg*POU2 | KRTSIETNIRVALEKSFQQNPKPSSEEITSVADQLSMEKEVVRVWFCNRRQKQKR | oyster | *C. gigas* | EKC20170.1 |
| *Hs*POU2 | KRTSIETNVRFALEKSFLANQKPTSEEILLIAEQLHMEKEVIRVWFCNRRQKEKR | human | *H. sapiens* | XP_011541043.1 |
| *Cg*POU4 | KRTSIAAPEKRSLEAYFAVQPRPSGEKIAQIAEKLDLKKNVVRVWFCNQRQKQKR | oyster | *C. gigas* | XP_011456596.1 |
| *Hs*POU4 | KRTSIAAPEKRSLEAYFAIQPRPSSEKIAAIAEKLDLKKNVVRVWFCNQRQKQKR | human | *H. sapiens* | NP_004566.2 |
| *CgPOU6* | KRKRRTSFTPQALEVLNNFFERNTHPSGAEMTELSDKLNYDREVIRVWFCNKRQA | oyster | *C. gigas* | EKC42925.1 |
| *Hs*POU6 | RRTSFTPQALEILNAHFEKNTHPSGQEMTEIAEKLNYDREVVRVWFCNKRQALKN | human | H. sapiens | NP_001159490.1 |
| *Ceh18* | KRTNLDMNQRNALDTFFALNPRPDHDKMTDIANSLELDRDVVRVWFCNRRQKMRR | nematode | *C. elegans* | AAA52203. |

#### Table S3

**A.**

| | | | |
|---|---|---|---|
| *Hs*ZFC1 | KHKCKYCSKVFGTDSSLQIHLRSHTGERPFVCSVCGHRFTTKGNLKVHFHRH----PQVKANPQL | KAI2595302.1 | Human |
| *Cg*ZFC1 | RHKCRLCQKVFGSDSALQIHIRSHTGERPFKCNICGNRFTTRGNLKVHFERHKAKYPHVKMNPHP | XP_034338577.1 | Oyster |

| | | | |
|---|---|---|---|
| *Hs*ZFC2 | ECLICHRVLSCQSSLKMHYRTHTGERPFQCKICGRAFSTKGNLKTHLGVHRTNTSIKTQHSCPI | KAI2595302.1 | Human |
| *Cg*ZFC2 | QCVICHRVLSCKSALQMHYRIHTGERPYKCKICGRAFTTKGNLKTHMGVHRMKPPIRMMHQCPV | XP_034338577.1 | Oyster |

**B.**

| | 919 922 | | | Bilateria |
|---|---|---|---|---|
| *Mm*SalL4 | CCTRCGKNFSSASALQIHERTHTGEKPFVCNICGRAFTTKGNLKVHYMTH | *M. musculus* | Q8BX22.2 | Mammalia |
| *Pp*SalL | VCNACGKNFSSASALQIHERTHTGEKPFACNVCGRAFTTKGNLKVHIGTH | *P. platessa* | CAB1440638.1 | Teleostei |
| *Ap*SalL | SCHVCGKQFQSDSALQIHMRTHTGEKPFKCHICGRAFTTKGNLKVHIGTH | *A. planci* | XP_022083928.1 | Echinodermata |
| *Dm*Spalt | FCHVCRRNFSSSSALQIHMRTHTGDKPFQCNVCQKAFTTKGNLKVHMGTH | *D. melanogaster* | NP_001245983.1 | Arthropodia |
| *Ce*SEM4 | QCGVCFKHFSSSSALQIHMRTHTGDKPFKCDMCGRAFTTRGNLKVHMGTH | *C. elegans* | CAA95798.1 | Nematode |
| *Oe*SalL | QCQVCQKGFSSASALQIHIRTHTGDKPFKCNVCGKAFTTKGNLKVHMGTH | *O. edulis* | XP_048751854.1 | Mollusca |
| *Cg*SalL | QCQVCQKGFSSASALQIHIRTHTGDKPFKCTVCGKAFTTKGNLKVHMGTH | *C. gigas* | XP_034338577.1 | Mollusca |
| *Hr*Sal | QCMTCMKTFSSASALQIHTRTHTGDKPFKCTVCGKAFTTKGNLKVHMGTH | *H. rubra* | XP_046576113.1 | Mollusca |
| *Pv*SalL | LCPTCHKTFSSGSALMIHNRTHTGDKPFKCNVCGKAFTTRGNLKVHMGTH | *P. vulgata* | XP_050398939.1 | Mollusca |
| *Of*SalL | LCQVCIKNFSSASALQIHMRTHTGDKPFKCTVCGKAFTTKGNLKVHMGTH | *O. fusiformis* | QRF78330.1 | Annelide |
| *Ml*Zfp | TCELCGKSCSCRSALEIHMRTHTKEKPFHCGQCGKGFSTKGNMKQHELTH | *M. lignano* | PAA52534.1 | Platyhelminthe |
| *Rs*Zfp | ICNVCSKNFSSASALQIHNRTHTGEKPYKCDVCGRAFTTKGNLKVHTGTH | *R. sordida* | CAF0925800.1 | Rotifera |
| *Ed*Zfp | FCNECGRSFTRSSTLVAHKRIHTGEKPFMCEICGRAFRQPGNLSRHRLTH | *E. diaphana* | KXJ20250.1 | Cnidaria |
| *Hc*Zfp | KCKHCGKEFGRSNNMKAHTRIHTGEKPFECRFCGKAFRFGNHLTIHERSH | *H. californensis* | AHA51381.1 | Ctenophora |
| *Ta*Zfp | KCKICGKAFSRPWLLQGHIRTHTGEKPFGCEKCNRSFADRSNLRAHMQTH | *T. adhaerens* | XP_002108251.1 | Placozoa |
| *Aq*Zfp12 | QCQFCKRVFTQASTLRSHERLHTGEKPYKCDHCGKAFTQSAGLRSHLKTH | *A. queenslandica* | XP_003387259.2 | Porifera |

Ecdyzoa
Lophotrochozoa
Deuterostomia
Protostomia
Epitheliozoa
Porifera
Metazoa

**Table S4**

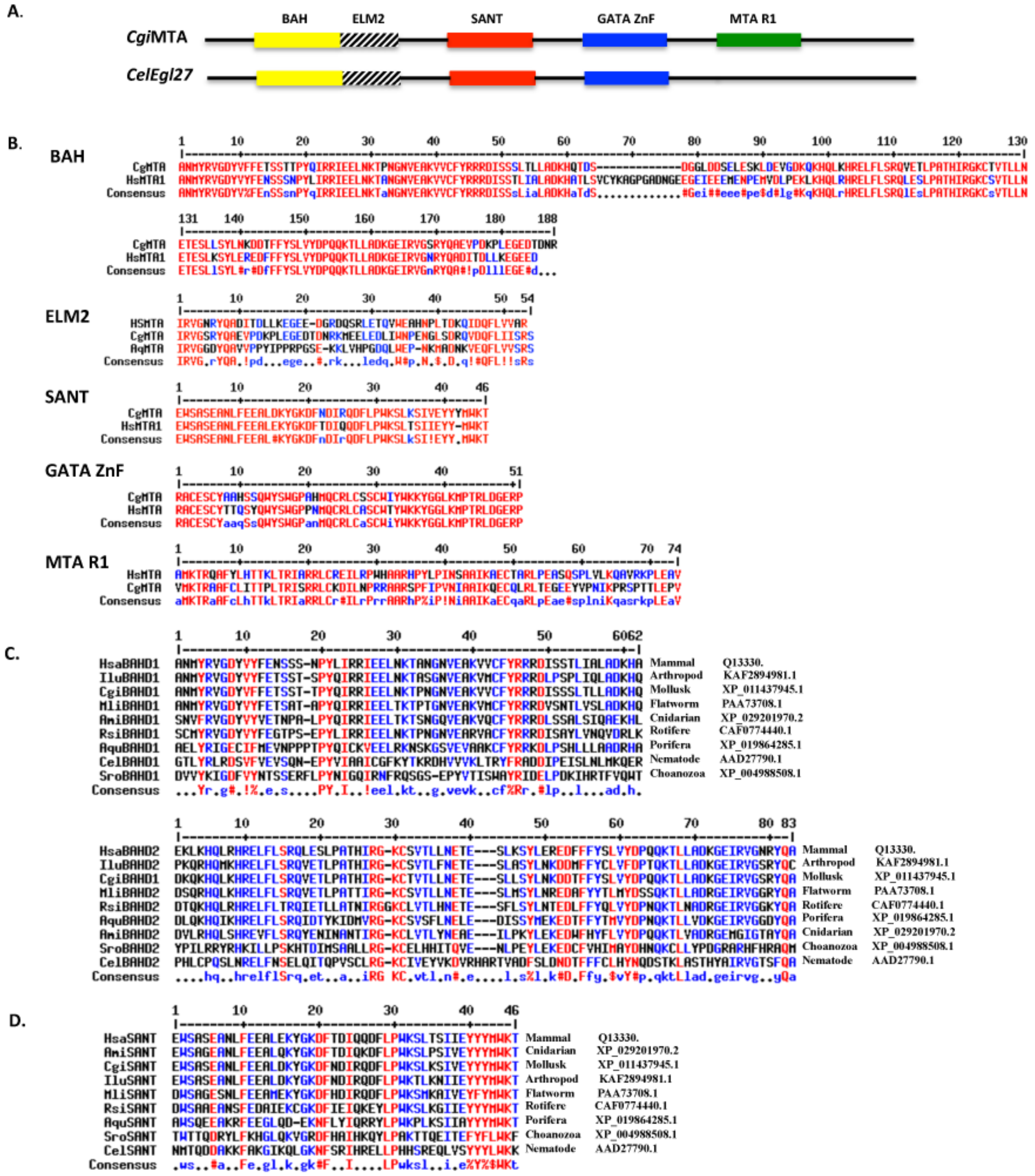
A.
CgiMTA
CelEgl27
BAH
ELM2
SANT
GATA ZnF
MTA R1
B.
BAH
ELM2
SANT
GATA ZnF
MTA R1
CgMTA
HsMTA1
HsMTA
AqMTA
Consensus
C.
HsaBAHD1 Mammal Q13330.
IluBAHD1 Arthropod KAF2894981.1
CgiBAHD1 Mollusk XP_011437945.1
MliBAHD1 Flatworm PAA73708.1
AmiBAHD1 Cnidarian XP_029201970.2
RsiBAHD1 Rotifere CAF0774440.1
AquBAHD1 Porifera XP_019864285.1
CelBAHD1 Nematode AAD27790.1
SroBAHD1 Choanozoa XP_004988508.1
HsaBAHD2 Mammal Q13330.
IluBAHD2 Arthropod KAF2894981.1
CgiBAHD1 Mollusk XP_011437945.1
MliBAHD2 Flatworm PAA73708.1
RsiBAHD2 Rotifere CAF0774440.1
AquBAHD2 Porifera XP_019864285.1
AmiBAHD2 Cnidarian XP_029201970.2
SroBAHD2 Choanozoa XP_004988508.1
CelBAHD2 Nematode AAD27790.1
D.
HsaSANT Mammal Q13330.
AmiSANT Cnidarian XP_029201970.2
CgiSANT Mollusk XP_011437945.1
IluSANT Arthropod KAF2894981.1
MliSANT Flatworm PAA73708.1
RsiSANT Rotifere CAF0774440.1
AquSANT Porifera XP_019864285.1
SroSANT Choanozoa XP_004988508.1
CelSANT Nematode AAD27790.1

#### Table S5

| | | | | | | |
|---|---|---|---|---|---|---|
| *Hs*, *Homo sapiens* (Mammalia) | Mammalia | | | | | |
| *Mm*, *Mus musculus* (Mammalia) | | | | | | |
| *Lp*, *Liparis tanakae* (snail fish) | | Gnathostoma | | | | |
| *Ol*, *Oryzias latipes* (medaka fish) | | | Chordata | | | |
| *Pp*, *Pleuronectes platessa* (plaice) | | | | | | |
| *Ar*, *Amblyraja radiata* (thorny skate) | Chondrychthyan | | | Deuterostoma | | |
| *Pm*, *Petromyzon marinus* (lamprey) | | Agnatha | | | | |
| *Bb*, *Branchiostoma belcheri* (Amphioxus) | Cephalochordata | | | | | |
| *Ac*, *Acanthaster planci* (star fish) | Echinodermata | | | | | |
| *Aj*, *Anneissia japonica* (Crinoidea) | | | | | | |
| *Dm*, *Drosophila melanogaster* (insect) | Arthropoda | | | | | |
| *Il*, *Ignelater luminosus* (insect) | | | | | | |
| *Cg*, *Crassostrea gigas* (Pacific oyster) | | | | | | |
| *Oe*, *Ostrea edulis* (European oyster) | Mollusca | | | | | |
| *Pv*, *Patella vulgata* (limpet) | | | | | Epitheliozoa | |
| *Hr*, *Haliotis rubra* (abalone) | | | | Protostoma | | Metazoa |
| *Of*, *Owenia fusiformis* (Annelid) | Annelidea | | | | | |
| *Sm*, *Schmidtea mediterranea* (Planaria) | Plathelminthes | | | | | |
| *Ml*, *Macrostomum lignano* (flat worm) | | | | | | |
| *Ce*, *Caenorhabditis elegans* (roun worm) | Nematoda | | | | | |
| *Rs*, *Rotaria sp. Silwood* (Rotifer) | Rotifera | | | | | |
| *Ha*, *Hydractinia echinata* (snail fur hydroid) | | | | | | |
| *Nv*, *Nematostella vectensis* (star anemone) | | | | | | |
| *Aa*, *Aurelia aurita* (blue jellyfish) | | | Cnidaria | | | |
| *Of*, *Orbicella faveolata* (star coral) | | | | | | |
| *Am*, *Acropora millepora* (elkhorn coral) | | | | | | |
| *Ml*, *Mnemiopsis leidyi* (Ctenary) | | | Ctenophora | | | |
| *Ta*, *Trichoplax adherens* (Placozoa) | | | Placozoa | | | |
| *Ol*. *Oscarella lobularis* (Homoscleromorpha) | | | | | | |
| *Sc*. *Sycon ciliatum* (Calcarea) | | | | | | |
| *Om*. *Oopsacas minuta* (Hexactinellida) | | | Porifera | | | |
| *Aq*, *Amphimedon queenslandica* (Demosponge) | | | | | | |
| *Hd*. *Halisarca dujardinii* (Demosponge) | | | | | | |
| *Sr*, *Salpingoeca rosetta* (Choanozoa) | | | Choanoflagellatea | | | Protozoa |

#### Table S6

### List of constructs used in this study

**For samples please contact:**
**For information please contact :**
***Cg* for *Crassostrea gigas***

#### Oyster Sox2

**Reference number**
10 pGEX 5X2-CgSox2
**GATEWAY**
pHGWA His6: + 2.8 kDa
pHGGWA His6 GST: + 28.3 kD a
pHXGWA His6 TRX: + 14.6 kDa
12 pDON221CgSox2
**Bacterial vectors**
13 pHGWA His6-CgSox2
14 pHGGWA His6-GST-CgSox2
15 pHxGWA His6-TRX-CgSox2
**Mammalian vectors**
16 pcDNA10 3xFlag- CgSox2
17 pcDNA3 MycS1- CgSox2
18 pcDNA5 FRT/SH/GW - CgSox2
**Vectors for biotinylation**
19 pBir CgSox2

21 pSBTet-StrepTag-HA-CgSox2 Addgene 60510
22 pSBTetPur Flag-APEX2-CgSox2 Addgene 60507

#### Oyster POU2

30 pBKS2- CgPOU2 X2
34 pCMSKz-CgPOU2X2
**GATEWAY**
35 pDON221 CgPOU2X2
**Bacterial vectors**
36 pHGWA His6-CgPOU2
37 pHGGWA His6-GST-CgPOU2
38 pHXGWA His6-TRX CgPOU2
**Mammalian vectors**
39 pCDNA10 3xFlag CgPOU2x2
40 pCDNA3 Myc S1 CgPOU2x2
41 pCDNA5/FRT/TO/SH/GW CgPOU2x2
**Sleeping Beauty**
42 pSBbiEF1a RB-HA-Tap-Tag-POU2 pSBTet 60522 Blasticidine
43 pSBbiEF1a RB-Myc-POU2 pSBTet 60522 Blasticidine
44 pSBTet-HA-Tap-TAG-POU2 pSBTet 60510 Blasticidine
45 pSBTet Puro POU2 pSBTet 60507 puromycine

#### Oyster POU3

**GATEWAY**
64 pDON221 CgPOU3
**Bacterial vectors**
65 pHGWA His6-CgPOU3
66 pHGGWA His6-GST-CgPOU3
67 pHXGWA His6-TRX CgPOU3
**Mammalian vectors**
68 pCDNA10 3xFlag CgPOU3
69 pCDNA3 Myc S1 CgPOU3
70 pCDNA5/FRT/TO/SH/GW CgPOU3
***Sleeping Beauty**
71 pSBTet HA-TAP-TAG-POU3 pSBTet 60510 Blasticidine
72 pSBbiEF1a HA-TAP-TAG-POU3 pSBTet 60522 Blasticidine
73 pSBbiEF1a Flag-POU3 pSBTet 60522 Blasticidine
74 pSBTet Puro Flag-POU3 pSBTet 60507 Puromycine

#### Oyster POU4

**GATEWAY**
92 pDON221 CgPOU4
**Bacterial vectors**
93 pHGWA His6-CgPOU4
94 pHGGWA His6-GST-CgPOU4
95 pHXGWA His6-TRX CgPOU4
**Mammalian vectors**
96 pCDNA10 3xFlag CgPOU4
97 pCDNA3 Myc S1 CgPOU4
98 pCDNA5/FRT/TO/SH/GW CgPOU4
**Sleeping Beauty**
99 pSBTet BR Myc-POU4 pSBTet 60510 Blasticidine
100 pSBTet BR Flag-POU4 pSBTet 60510 Blasticidine

### Oyster POU6

**GATEWAY**
116 pDON221 CgPOU6
**Bacterial vectors**
117 pHGWA His6-CgPOU6
118 pHGGWA His6-GST-CgPOU6
119 pHXGWA His6-TRX CgPOU6
**Mammalian vectors**
120 pCDNA10 3xFlag CgPOU6
121 pCDNA3 Myc S1 CgPOU6
122 pCDNA5/FRT/TO/SH/GW CgPOU6
**Sleeping Beauty**
123 pSBTet Myc-POU6 pSBTet 60510 Blasticidine
124 pSBTet Flag-POU6 pSBTet 60510 Blasticidine
125 pSBbi EF1a RB-Myc-POU6 pSBTet 60522 Blasticidine
126 pSBTet Puro Flag-POU6 pSBTet 60522 Blasticidine

### Oyster MTA

132 pBKS2CgMTA3 X1
133 pBKS2CgMTA3 X2
**Mammalian vectors**
134 pEGFPC1 CgMTA X1
136 pDsRedC1 CgMTA X1
138 pCS3MT-CgMTA X1
140 pBirSumo CgMTA X1
**Sleeping Beauty**
142 pSBTet-StrepTag-Myc-CgMTA pSBTet 60510 Blasticidine
143 pSBTet Puro Myc-MTA pSBTet 60507 Puromycine
144 pSBTet Hygro mRFP-MTA pSBTet 60508 Hygromycine
145 pSBTet Hygro Myc-MTA pSBTet 60508 Hygromycine

### Oyster SalL

151 pGEX4T2 CgSal Cter MW 66,55 kDa pI: 8,30 : GST + the last 360 AAs of SalL
157 pBKS2 CgSal X1 Full Length
158 pCMVBirS CgSalX1
160 pCMVBirS CgSalX1-PolyA
163 pCMV BirFlag CgSalX1 PolyA
**Sleeping Beauty**
167 pSBTet Streptag-Flag-CgSal pSBTet 60510 Blasticidine
168 pSBTet Puro Flag-SalL4 pSBTet 60507 Puromycine

### Oyster Lin28

170 pCS3MycCgLin28 X3
**Sleeping Beauty**
173 pSBTetBR StrepTag-Myc-CgLin28X3 pSBTet 60510 Blasticidine

### Xenopus Histone H2B

200 pSBTet HcRed-XH2B pSBTet 60510 Blasticidine

*pSBtet vectors were a gift from Eric Kowarz (Addgene plasmid # 60507; #60508; #60510 http://n2t.net/addgene:60507; #60508; #60510; RRID: Addgene_60507 ; _60508; _60510) (Kowarz et al., 2015)
